# Invasion of glioma cells through confined space requires membrane tension regulation and mechano-electrical coupling via Plexin-B2

**DOI:** 10.1101/2024.01.02.573660

**Authors:** Chrystian Junqueira Alves, Theodore Hannah, Sita Sadia, Christy Kolsteeg, Angela Dixon, Robert J. Wiener, Ha Nguyen, Murray J. Tipping, Júlia Silva Ladeira, Paula Fernandes da Costa Franklin, Nathália de Paula Dutra de Nigro, Rodrigo Alves Dias, Priscila V. Zabala Capriles, José P. Rodrigues Furtado de Mendonça, Paul Slesinger, Kevin Costa, Hongyan Zou, Roland H. Friedel

**Author notes:** These authors jointly supervised this work (Chrystian Junqueira Alves, Hongyan Zou, Roland H. Friedel). **Corresponding authors**: Chrystian Junqueira Alves; Hongyan Zou; Roland Friedel.

## Abstract

Glioblastoma (GBM) is a malignant brain tumor with uncontrolled invasive growth. Here, we demonstrate how GBM cells usurp guidance receptor Plexin-B2 to gain biomechanical plasticity for polarized migration through confined space. Using live-cell imaging to track GBM cells negotiating microchannels, we reveal active endocytosis at cell front and filamentous actin assembly at rear to propel GBM cells through constrictions. These two processes are interconnected and governed by Plexin-B2 that orchestrates cortical actin and membrane tension, shown by biomechanical assays. Molecular dynamics simulations predict that balanced membrane and actin tension are required for optimal migratory velocity and consistency. Furthermore, Plexin-B2 mechanosensitive function requires a bendable extracellular ring structure and affects membrane internalization, permeability, phospholipid composition, as well as inner membrane surface charge. Together, our studies unveil a key element of membrane tension and mechanoelectrical coupling via Plexin-B2 that enables GBM cells to adapt to physical constraints and achieve polarized confined migration.

## INTRODUCTION

Cancer is not only a disease of uncontrolled proliferation but also uncontrolled infiltration. This is particularly pertinent for glioblastoma (GBM), the most common and lethal brain cancer that is notorious for wide dissemination in the brain ^1–3^. Invading GBM cells frequently experience physical constraints from confined interstitial space, they need to sense and respond to mechanical forces to rapidly adjust migratory behavior. Earlier research has investigated molecular drivers of cell intrinsic motility (e.g., mesenchymal shift) ^4^, but how GBM cells respond to extrinsic factors is less understood.

Physical constraints result in nuclear deformation and cell body distortion ^5^, as well as local membrane stretching or ruffling. In intracranial transplants using patient-derived glioma stem cells (GSCs), we have detected large numbers of GBM cells invading along white matter tracts in the corpus collosum, with nuclei and cell body transformed into elongated fusiform shape in the direction of migration ^6^. Intriguingly, in response to constricted space, tumor cells can acquire a polarized migratory phenotype to rapidly escape crowded tissue regions ^5^; recent evidence also highlights polarization of lipids and inner membrane surface charge during cell migration ^7^. However, how cell mechanics is orchestrated with membrane electric charges that enable GBM cells to rapidly respond to physical constraints to achieve polarized migration through confined space remains unclear.

As tumor cells frequently usurp developmental pathways during malignant transformation, we have been focusing on axon guidance molecules in GBM invasion, particularly Plexin-B2, a membrane receptor initially cloned as a gene upregulated in GBM ^8^ and whose expression level correlates with poor survival of GBM patients ^9^. Notably, Plexin-B2 deletion can impede GBM invasion in patient-derived xenotransplant (PDX) models ^6^, however the biomechanical mechanisms remain opaque and Plexin-B2 function in confined migration has not been studied.

Aside from axon pathfinding, plexins also regulate a diverse range cell interactions, including immune activation, bone homeostasis, vascular remodeling, and cancer progression ^10^. Evolutionary studies revealed that plexins emerged more than 600 million years ago in a shared unicellular ancestor of metazoa and choanoflagellates, well before the appearance of nervous system ^11,12^. Plexins are transmembrane receptors with a highly conserved ring-shaped extracellular domain and an intracellular GAP (GTPase activating protein) domain that signals to cytoskeletons via deactivation of Ras/Rap small GTPases ^13,14^. While canonical plexin signaling occurs through dimerization upon binding to semaphorin ligands (exemplified in growth cone collapse) ^15,16^, recent evidence also implicated a mechanosensory function, e.g., Plexin-D1 in endothelial cells to detect blood flow sheer stress ^17^ and Plexin-B1 and -B2 in epidermal stem cells in response to compression forces ^18^. In echo, our own work has shown that Plexin-B2 controls actomyosin tension and cell stiffness of human embryonic stem cells (hESCs) and neural progenitor cells (hNPCs), which is crucial for multicellular organization in hESC colony formation and neurodifferentiation ^19^.

Here, we investigated cell mechanics, in particular, membrane tension in connection with membrane electrical features during confined migration. By applying advanced fluorescent dyes and protein probes, combined with live cell imaging, we tracked the spontaneous migration of GBM cells into 3D microchannels containing constrictions, mimicking the mechanical stress experienced by invading GBM cells. We unveiled regionalized membrane internalization/endocytosis at cell front and filamentous (F) actin assembly at cell rear during confined migration, two interconnected processes that are linked to asymmetry of phospholipid composition, inner membrane surface charges, and calcium accumulation. These processes were governed by Plexin-B2, signaling through a bendable extracellular ring, leading to polarized migration through confined space. Together, our results underscore the importance of Plexin-B2-regulated membrane tension and mechanoelectrical coupling for confined migration. Targeting these interconnected processes may provide a new angle to tackle GBM invasion.

## RESULTS

### Passage through confined space enhances migratory momentum and consistency of GBM cells

To gauge the capability of GBM cells for confined migration, we conducted invasion assays with patient-derived GBM stem cell (GSC) lines using microdevices containing microchannels (12 µm wide, 10 µm high) with periodic 3 or 8 µm constrictions (**Fig. 1A**). The microchannels were laminin-coated with no chemoattractants applied, thus migration of GSCs into the microchannels was spontaneous, driven by intrinsic motility. To visualize cell movement, we labeled GSCs with live cell nuclear dye NucSpot before being seeded in inflow ports, and tracked cell movement by time-lapse live cell imaging for up to 21 hr. To ascertain if confined migration might be impacted by molecular subtypes of GBM, we tested GSC lines of either mesenchymal (SD2) or proneural (SD3) subtype ^20^, which both had shown wide dissemination in intracranial transplants, with nuclei and cell bodies transformed into fusiform shapes aligned along white matter tracts ^6^.

**Figure 1.**
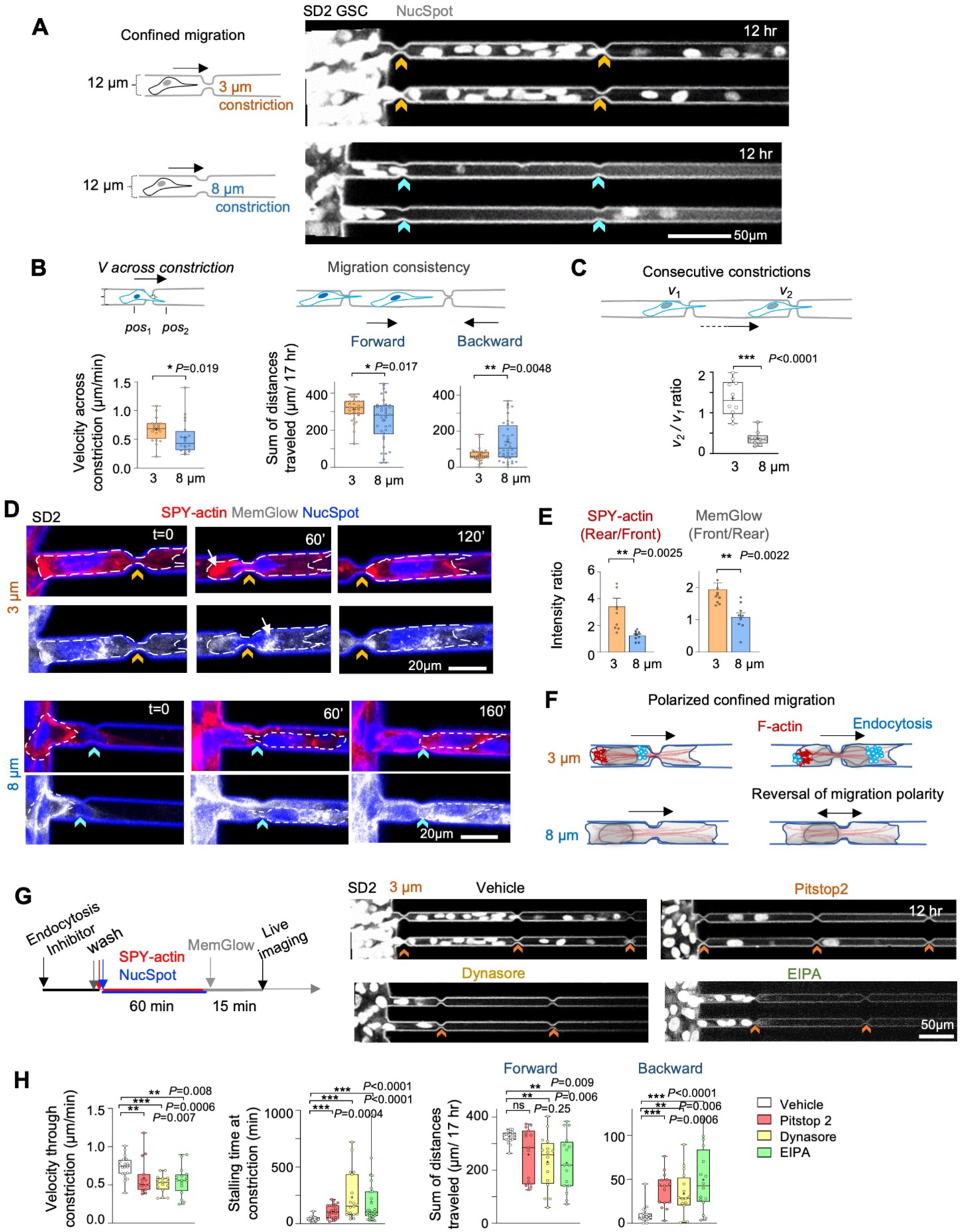
Confined migration of GSCs requires endocytosis. (A) Left, schematic illustration of spontaneous migration in microchannels with periodic 3 or 8 µm constrictions. Right, still frames from videography show SD2 GSCs labeled with NucSpot invading into microchannels at 12 hr after seeding. Chevrons point to constrictions. (B) Quantifications (diagram on top) of velocity across constrictions (n=20 cells per condition) and sums of distances traveled by forward or backward movements over 17 hr (n=31-40 cells per condition). Box plots show 25–75% percentiles, minimal and maximal values (whiskers), median (line), and mean (cross). Mann–Whitney– Wilcoxon test. (C) Box plots show velocity ratio when crossing 2nd vs. 1st constriction. n=10 cells per group. Two-sided unpaired t-test. (D) Still frames from live-cell fluorescent imaging of SD2 GSCs traversing constrictions (note different time frames for 3 or 8 µm). Dashed lines outline cell boundary. Arrows points to F-actin at rear zone (SPY555-actin) or MemGlow-labeled endosomes at front zone of cell. (E) Quantifications of fluorescence intensity ratio for SPY-actin or MemGlow at rear vs. front zones during confined migration through 3 or 8 µm constrictions. n=10 cells per group. Two-sided unpaired t-test. (F) Schematic illustration of polarized migration through constrictions, which further augments migration consistency. Note regionalized endocytosis at cell front and F-actin assembly at rear to propel cells through constriction, more efficient for 3 µm than 8 µm constrictions. (G) Left, experimental timeline for endocytosis inhibitor treatment before microchannel invasion assay. Right, still frames from videography show stalled migration of inhibitor treated SD2 GSCs at 12 hr after seeding. (H) Box plots of velocity through constriction, stalling time at constrictions, and sum of forward and backward movements affected by endocytosis inhibitors. n=15 to 28 cells per condition. Kruskal–Wallis test followed by Dunn’s multiple comparisons test.

Remarkably, contrary to the assumption that narrower constrictions may slow down migration due to increased mechanical challenge, we observed that GSCs traversed 3 µm constrictions more efficiently than 8 µm ones, evidenced by more cells invading into the microchannels and faster speed (**Fig. 1A, B; S1A, B; Movie S1**). Moreover, passage through 3 µm constrictions further enhanced migration consistency (directional polarity), reflected by higher sums of forward movement than backward movement (**Fig. 1B; Movie S1**). These findings applied to both SD2 and SD3 GSC lines, with SD2 traversing the 3 µm constrictions at slightly lower velocity than SD3 (∼0.7 vs. 1 µm/min) (**Movie S2;** see also **Fig. 4B**).

When following the same GSC passing through two consecutive 3 µm constrictions, passage through the first increased the velocity for the second (by ∼25%) (**Fig. 1C**). This again applied to both SD2 and SD3 (**Fig. S1C, D**). In contrast, GSC passing through two consecutive 8 µm constrictions displayed lower momentum at the second passage (**Fig. 1C; S1C, D; Movie S1**). By contrast, hNPCs derived from hESCs displayed much lower migratory capacity to negotiate microchannels (**Fig. S1E, S1F Movie S3**), indicating a high cell type-specific biomechanical plasticity in GSCs for spontaneous migration through confined space.

### Confined migration involves membrane internalization via endocytosis

The adaptation to physical constraints requires dynamic reorganization of both cytoskeleton and plasma membrane. We thus applied live cell fluorogenic dyes SPY-actin and MemGlow, revealing robust F-actin assembly at cell rear and numerous MemGlow-labeled endocytic vesicles at the front zone, more evident in GSCs traversing 3 µm than 8 µm constrictions (**Fig. 1D**). There was also a smaller proportion of vesicles gathered at the rear zone in constricted cells, particularly after passage, while NucSpot labeling highlighted nuclear deformation of constricted cells (**Fig. 1A, D**). Hence, physical constraint triggers not only regionalized F-actin assembly at rear zone to propel cells through constrictions, but also active membrane internalization at cell front (**Fig. 1E**).

The balance of endocytosis and exocytosis is a central process to adjust local membrane tension ^21–23^. We thus tested if endocytosis is required for confined migration. We treated GSCs with three small molecule inhibitors targeting different pathways of endocytosis: PitStop 2 (a cell permeable clathrin inhibitor), Dynasore (a noncompetitive inhibitor of dynamin), and EIPA (an inhibitor of Na^+^/H^+^ exchange pump that blocks macropinocytosis by lowering pH near plasma membrane and thereby inhibiting actin remodeling via Rho) ^24–27^. All three inhibitors disrupted accumulation of MemGlow-labeled endocytic vesicles at cell front, but notably also actin network at cell rear (**Fig. 1G; Fig. S2A, B**). This illustrated that the two processes are interconnected, required for confined migration as measured by velocity through constrictions, stalling time, and migration consistence/polarity in microchannels (**Fig. 1G, H; Movie S4**). By comparison, in 2D cultures of GSCs, endocytosis inhibitors had no major impact on F-actin (phalloidin), proliferation (Ki67 IF), or migration (wound closure assay; except for a modest delay by Dynasore at 24-72 hr and EIPA at 72 hr) (**Fig. S2C, D**).

### Guidance receptor Plexin-B2 controls cortical actin and membrane tension of GSCs

So far, our results showed polarized endocytosis at front and F-actin at rear during confined migration. Our next question pertained to molecules orchestrating these two interconnected processes. Guidance receptor Plexin-B2 is an attractive candidate, as it is a transmembrane protein controlling actin contractility/cell stiffness in hESCs and hNPCs ^19^, and it is upregulated in GBM and promotes invasion via modulating actomyosin and cell adhesion ^6,9^, but its role in confined migration is unknown.

We generated GSCs with CRISPR/Cas9-mediated *PLXNB2* knockout (PB2 KO), or lentiviral overexpression (PB2 OE), confirmed by Western blot and IF staining (**Fig. 2A-C**). Peculiarly, we observed elongated cell processes of PB2 KO cells in 2D culture (**Fig. S3A**), a phenotype better visualized by live cell imaging of GSCs labeled with plasma membrane dye NR12A ^28^ (**Fig. S3B; Movie S5**). This could be a sign of reduced membrane dynamics to withdraw extended cell processes in the absence of Plexin-B2. Consistently, 3D surface rendering of NR12A-labeled GSCs revealed a more ruffled plasma membrane of PB2 KO cells (**Fig. S3C**).

**Figure 2.**
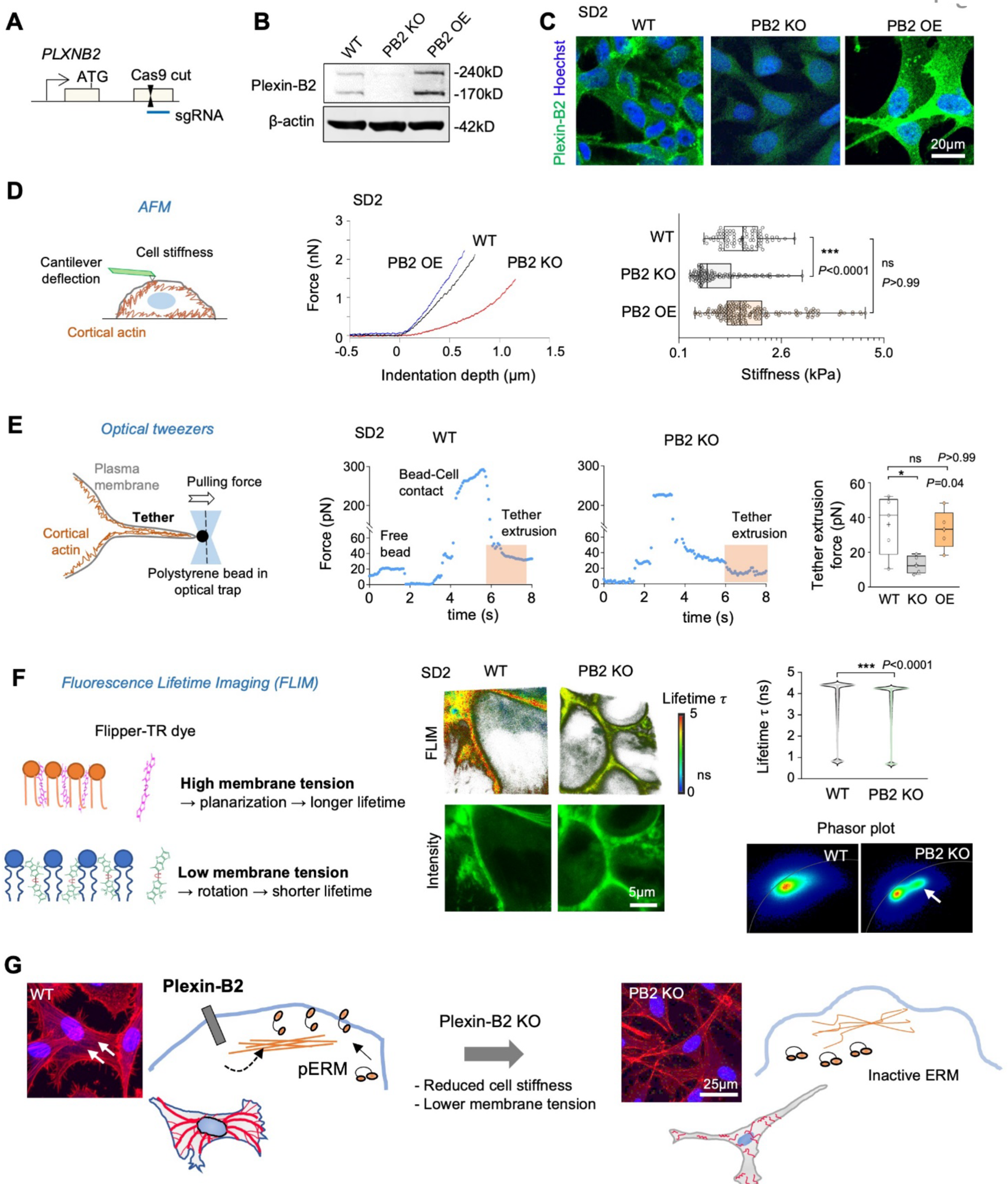
Plexin-B2 regulates cytoskeletal and membrane tension in GSCs. (A) Schematic of CRISPR/Cas9-mediated *PLXNB2* knockout (KO) with small guide (sg) RNA targeting second coding exon. (B) Western blots show Plexin-B2 expression in different SD2 GSCs, with β-actin as loading control. Note Plexin-B2 precursor at 240 kDa and mature form at 170 kDa. (C) IF images show Plexin-B2 expression in different SD2 GSCs, with Hoechst nuclear counterstain. (D) Left, schematic of atomic force microscopy (AFM) indentation method to probe cell stiffness by cantilever deflection. Middle, AFM indentation curves of different SD2 GSCs; right, box plots of cell stiffness, showing 25– 75% quartiles, median (line), and mean (plus sign). n= 6 cells per group. Kruskal–Wallis test followed by Dunn’s multiple comparisons test. (E) Left, depiction of membrane tension measurement with optical tweezers. Middle, force measurements during tether extrusion (shaded box). Right, quantifications of tether extrusion forces. n=5 cells per group. Kruskal–Wallis test followed by Dunn’s multiple comparisons test. (F) Left, schematic of FLIM of cell membranes labeled with Flipper-TR membrane dye, with low and high membrane tension associated with shorter and longer lifetimes, respectively. Middle top, representative FLIM images, with lifetime heatmap shown on right. Middle bottom, images show similar fluorescence intensities of Flipper-TR dye in WT and PB2 KO cells. Right top, violin plots show fluorescence lifetime from 3 images per group. Two-sided unpaired t-test. Right bottom, phasor plots of FLIM image data, with arrow indicating a shift to shorter lifetime values for PB2 KO cells. (G) Model of Plexin-B2 regulation of cortical contractility and membrane tension. Phalloidin staining show differences of F-actin network in WT and PB2 KO SD2 GSCs. DAPI for nuclear staining. Arrows point to stress fibers and spread-out contours of the WT GSCs.

We next performed micro-indentation measurements with atomic force microscopy (AFM) to directly gauge cell stiffness, a readout of cortical actin contractility (**Fig. 2D**). This revealed a lower stiffness for PB2 KO cells (mean ∼1 kPa) than WT cells (∼1.6 kPa), whereas PB2 OE cells displayed higher stiffness (∼1.8 kPa) (**Fig. 2D; S3D**). Hence, as in hESCs and hNPCs ^19^, Plexin-B2 appears to control also cortical actin contractility of hGSCs.

To measure membrane tension, we applied optical tweezers to determine the force required to pull out plasma membrane tethers, which was lower for PB2 KO cells (mean ∼12.8 pN) than WT (∼36 pN) or PB2 OE SD2 cells (∼33 pN) (**Fig. 2E**). Interestingly, unlike sustained PB2 OE, acute overexpression of PB2 in a doxycycline-inducible fashion also led to reduced tether pulling force (**Fig. S3E, F**), illustrating that acute imbalance of cortical contractility can perturb membrane tension. In support, pharmacological inhibition of myosin contractility (blebbistatin) or actin assembly (latrunculin) markedly reduced the tether pulling forces for WT GSCs (phenocopying PB2 KO), with no further effect on PB2 KO cells (low baseline actomyosin contractility) (**Fig. S3G**).

As a second way to gauge plasma membrane tension in live cells, we utilized Flipper-TR, a membrane dye that changes fluorescence lifetime according to membrane tension ^29^. Fluorescence lifetime imaging (FLIM) revealed shorter Flipper-TR lifetimes in PB2 KO cells (indicating lower membrane tension) for both SD2 and SD3, as well as GSCs with Dox-inducible shRNA knockdown of PB2 (**Fig. 2F; S4A, B**). Echoing optical tweezers results, sustained PB2 OE did not alter Flipper-TR lifetimes, but acute PB2 OE (6 hr after Dox induction) did, which was recovered by 24 or 48 hr (**Fig. S4A, C**).

To understand the connection of cortical actin and membrane tension, we examined phosphorylated ezrin-radixin-moesin (ERM), which can act as a linker between cortical actin and membrane proteins ^30^. IF staining revealed co-localization of Plexin-B2, pERM, and F-actin near cell membrane of WT GSCs, a patterned perturbed by PB2 KO (reduced pERM) or PB2 OE (aberrant pERM aggregation, particularly in SD3 cells) (**Fig. S5A, C**). The levels of pERM were only modestly decreased by PB2 KO shown by WB, while total ezrin levels were comparable (**Fig. S5B, D**). In sum, biomechanical measurements by AFM, optical tweezers, and Flipper-TR FLIM support the model that Plexin-B2 mediates cortical actin and membrane tension in GSCs, and IF staining indicate that this process is mediated via regulation of pERM (**Fig. 2G**).

### Plexin-B2 function impacts endocytosis and membrane internalization of GSCs

As cell membrane tension may affect endocytosis and membrane internalization, we investigated next the impact of Plexin-B2 on endocytosis by performing a dextran uptake assay, a standard approach that has also been applied previously to demonstrate endocytosis in growth cone collapse stimulated by semaphorins ^15,16^. Exposure to dextran (MW 10 kDa, conjugated to Alexa488) for 40 min indeed resulted in intracellular fluorescent puncta in WT GSCs, indicating endosomal uptake, but the number of endosomal puncta was significantly lower in PB2 KO cells (**Fig. 3A**).

**Figure 3.**
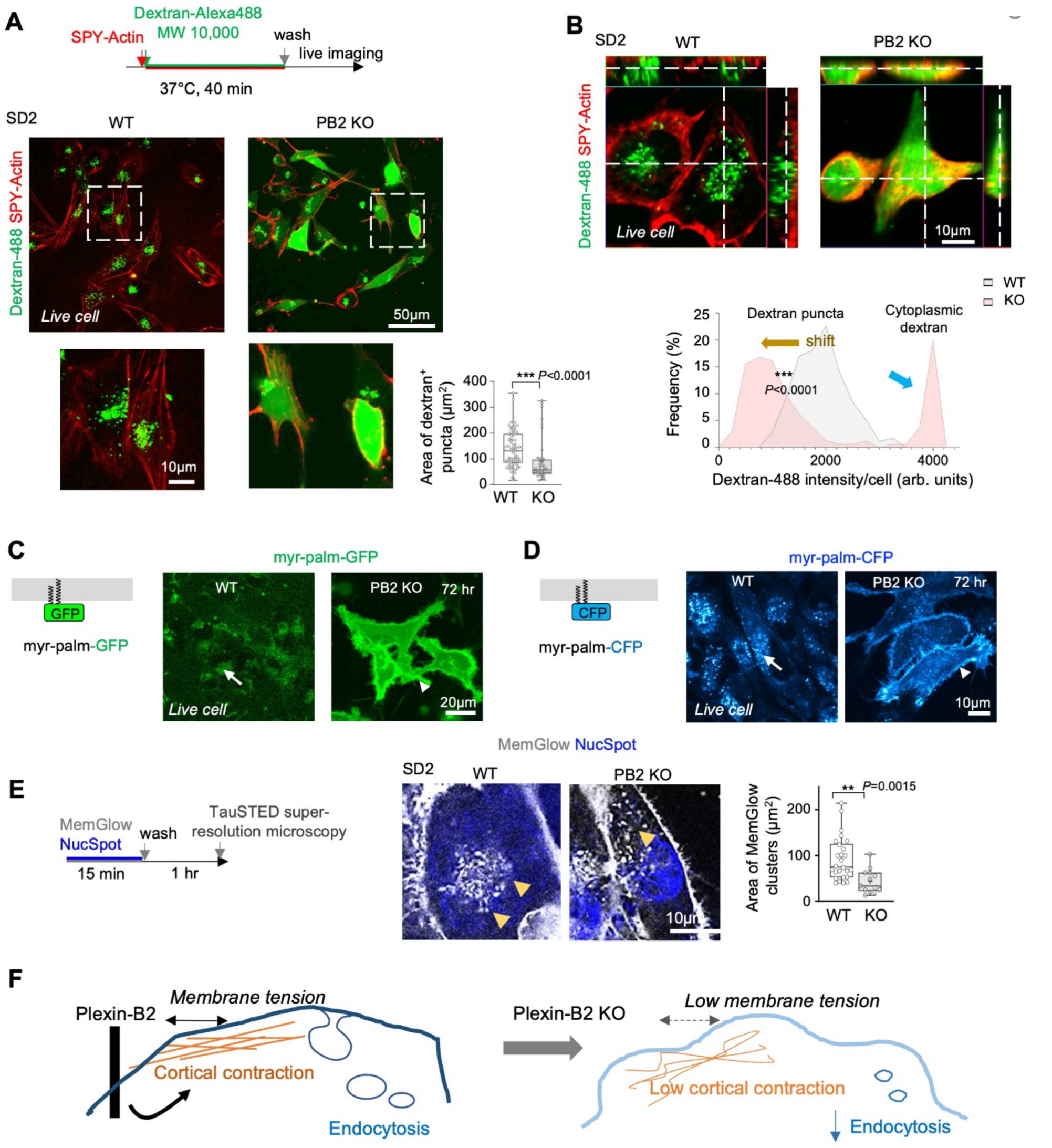
Plexin-B2 affects membrane internalization in GSCs. (A) Top, timeline for dextran uptake assay. Bottom, live-cell imaging of WT and PB2 KO SD2 GSCs labeled with SPY-Actin and exposed to dextran-Alexa488. Enlarged images of boxed areas are shown below. Quantification of the areas of dextran^+^ clusters per cell are shown in box plots, with 25–75% quartiles, median (line), and mean (plus sign). n=85 cells for WT, n=44 cells for PB2 KO. Mann–Whitney–Wilcoxon test. (B) Top, live cell confocal plane images of WT and PB2 KO GSCs with side views of z-stacks showing intracellular localization of diffuse dextran-Alexa 488 signals in PB2 KO cells in addition to dextran endosome signals. In contrast, WT cells contained only dextran^+^ endosomes. Bottom, histograms show fluorescence profiles showing bimodal distribution of dextran-Alexa 488 fluorescence intensities in PB2 KO GSCs (blue and brown arrows). n=177 cells for WT, n=161 cells for PB2 KO. Mann–Whitney–Wilcoxon test. (C, D) Left, schematic of myr-palm-GFP or -CFP attached to inner membrane leaflet. Right, live cell fluorescence imaging at 72 hr after transfection shows internalization of myr-palm-GFP or -CPF on endomembranes (arrow) in WT GSCs, in contrast to membrane retention of the probes (arrowhead) in PB2 KO GSCs. (E) Left, schematic of TauSTED super-resolution microscopy of GSCs labeled with MemGlow. Middle, TauSTED live-cell images show reduced endosomes (arrowheads) in PB2 KO cells compared to WT. Right, box plots show areas of MemGlow clusters in each cell. n=26 cells for WT, n=13 cells for PB2 KO. Two-sided unpaired t-test. (F) Working model of regulation of cortical and membrane tension by Plexin-B2, affecting endocytosis and membrane permeability in GSCs.

Strikingly, in many PB2 KO cells we also observed bright uniform dextran fluorescent signals localized in the cytoplasm, as confirmed by confocal z-stack imaging (**Fig. 3A, B**). This suggested direct translocation of dextran through a leaky cell membrane. A fluorescence intensity histogram revealed two populations of dextran-containing KO cells, one containing fluorescent puncta, and one containing additionally also bright cytoplasmic signals (**Fig. 3B**). Similar results were also obtained with SD3 GSCs (**Fig. S6A**). PB2 OE led to a modest increase in dextran puncta, but also bright cytoplasmic fluorescence in some cells (**Fig. S6A**), indicating the impact of actomyosin hypercontractility on membrane permeability. Co-labeling of GSCs with lysotracker dye revealed a strong overlap of lysosomes with dextran puncta, but not with the diffuse cytoplastic dextran fluorescence observed in PB2 KO or OE cells (**Fig. S6B)**.

Both actomyosin inhibition by blebbistatin or endocytosis inhibitors led to increased cytoplastic dextran uptake in WT cells, mirroring the PB2 KO phenotype (**Fig. S6C; S7A-C**), whereas blebbistatin treatment attenuated the effect of PB2 OE in this regard (**Fig. S6C**). In further support of membrane leakage in PB2 KO or OE cells, cooling cells to 4 °C, which reduces membrane permeability by increasing phospholipid packing and membrane rigidity ^31^, markedly reduced cytoplasmic dextran fluorescence (**Fig. S6D**).

To further characterize membrane turnover in dependence of Plexin-B2, we introduced the membrane anchored fluorescent proteins myr-palm-GFP and -CFP (**Fig. 3C, D**). Live-cell imaging at 72 hr after transfection revealed that myr-palm-GFP or -CFP were mostly localized on intracellular vesicles in WT cells but retained on plasma membrane of PB2 KO cells (**Fig. 3C, D**). Super-resolution microscopy further demonstrated reduced MemGlow^+^ intracellular vesicles in PB2 KO cells (**Fig. 3E**). Likewise, FLIM imaging after incubation with Flipper-TR dye for 2 or 5 hr revealed reduced amount of internalized Flipper-TR dye in PB2 KO GSCs (**Fig. S6E)**. Altogether, Plexin-B2 manipulations affected endocytosis/membrane internalization and membrane permeability (**Fig. 3F**).

### Plexin-B2-deficient GSCs display compromised confined migration

Given that membrane tension/turnover were reduced in PB2 KO GSCs, we next investigated their capability for confined migration. Live cell videography revealed that significantly fewer PB2 KO SD2 or SD3 GSCs invaded into microchannels with 3 µm constrictions, with decreased velocity at constriction points, longer stalling times, and lower migration consistency (increased backward movements) (**Fig. 4A, B; Fig. S8; Movies S6, S7**). F-actin assembly at cell rear and endosomes at cell front were also disrupted in PB2 KO cells (**Fig. 4C**). PB2 OE cells performed better than PB2 KO cells, but worse than WT cells, with disorganized F-actin and endosomes (**Fig. S8B-F**).

**Figure 4.**
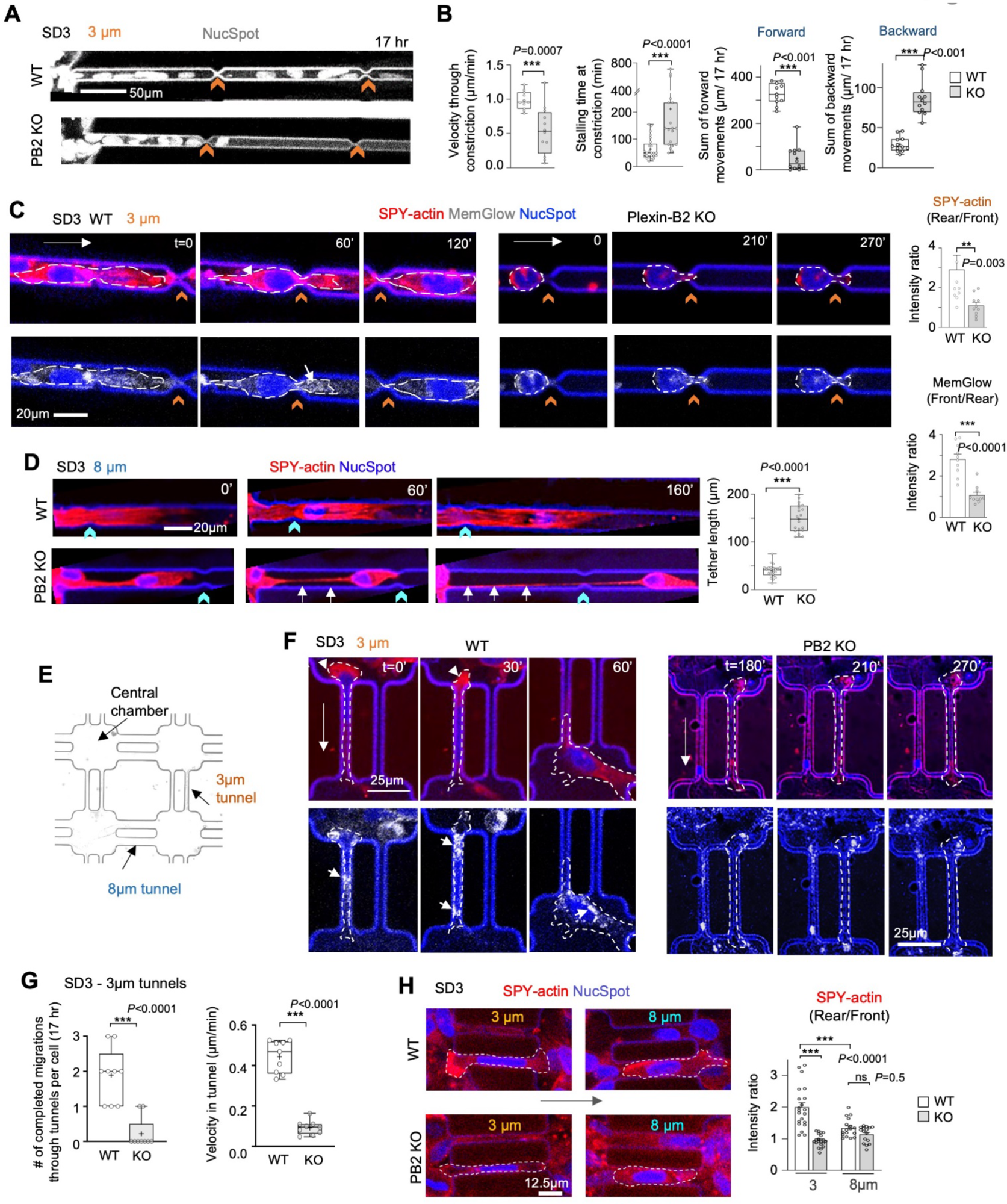
Plexin-B2 is required for migration of GSCs through confined space. (A) Still frames from live-cell videography show compromised migration of PB2 KO SD3 GSCs (visualized by NucSpot) into microchannels with 3 µm constrictions (denoted by orange chevron signs) as compared to WT cells. (B) Quantifications of velocity through constrictions (n=11-12 cells per genotype), stalling time at constrictions (n=19-25 cells), sums of distances traveled forward or backward over 17 hr (n=12 cells). Box plots show 25–75% quantiles, minimal and maximal values (whiskers), median (line), and mean (cross). Two-sided unpaired t-test. (C) Left, still frames from live-cell imaging of SD3 GSCs traversing 3 µm constrictions reveal F-actin assembly at rear zone (SPY555-actin, arrowhead) and MemGlow-labeled endosomes at front (arrow) in WT but not KO cells. Note different time stamps for WT and PB2 KO. Dashed lines delineate cell boundary. Right, quantifications of the ratio of fluorescence intensity for SPY-actin or MemGlow at rear vs. front zones during confined migration. n=10 cells per group. Mann–Whitney–Wilcoxon test. (D) Left, still frames from live-cell imaging of SD3 GSCs show a long tether (arrows) connecting PB2 KO cells in microchannel with 8 µm constriction. Right, box plot quantifications of SPY-actin tether length. n=24 cells for WT, n=20 cells for PB2 KO. Two-sided unpaired t-test. (E) Schematic of a microdevice with central chambers connected by pairs of narrow long exit tunnels of 50 μm length and 3 or 8 µm width. (F) Representative still images show successful passage of WT SD3 GSC through long narrow tunnel (3 µm) in 60 min timeframe, but stalled PB2 KO cell even after 270 min. Long arrows on left denote migration direction. Note F-actin (SPY-actin) at rear zone (arrowhead) and diffuse MemGlow-labeled endosomes (arrow) in WT, both reduced in PB2 KO GSC. Dashed lines delineate cell boundary. (G) Box plots show the number of successful migration events per cell through the 3 µm tunnel and the speed in the tunnel. n=9 cells per group. Two-sided unpaired t-test. (H) Left, still images show F-actin assembly at rear zone of WT GSCs traversing through tunnels, more prominent in 3 µm than 8 µm tunnels, and reduced in PB2 KO cells. Arrows denote direction of migration. Right, bar graphs show ratios of SPY-actin fluorescence intensities at rear vs. front during GSC passage through tunnels. For 3 µm, n=20-21 cells, for 8 µm, n=16-17 cells. One-way ANOVA followed by Tukey’s multiple comparison test. Data represent mean ± SEM.

Also of interest was a striking observation of long actin-containing tethers connecting PB2 KO GSCs in microchannels, reflective of failed dissociation from neighboring cells (**Fig. 4D; Movie S8**), in line with increased adhesiveness o GBM cells in the absence of Plexin-B2 ^6^.

To further characterize biomechanical processes during confined migration, we designed a microdevice with central chambers connected by pairs of tunnels of 3 or 8 µm width and 50 μm length (**Fig. 4E**). This design allows GBM cells to choose alternative exit routes, while the long micro tunnel presents continuous mechanical challenge. Live-cell imaging showed that WT GSCs were adept at exploring both 3 or 8 µm outlets and passing through tunnels; after passage, cells then altered direction in chambers and continued migration through the same or neighboring tunnels (**Fig. 4F; Movie S9**). In contrast, PB2 KO cells frequently stalled in tunnels, with reduced speed, less successful passages, and lower propensity to explore exit outlets or alter direction in central chambers (**Fig. 4F, G; S9A; Movie S9**).

For WT GSCs traversing micro tunnels, live-cell fluorescence imaging also captured F-actin assembly at rear zone and endosomes at cell front, more pronounced in 3 µm than 8 µm tunnels, but less evident in entrapped PB2 KO cells (**Fig. 4F, H**). Notably, the nuclei of constricted GSCs in micro tunnels were compressed into elongated shapes for both WT and PB2 KO (**Fig. 4F, H**), indicating that nuclear mechanical plasticity in response to physical constraints is still operational in the absence of Plexin-B2.

Recent studies showed that exposure to high fluid viscosity can increase cytoskeletal contractility through focal adhesions, leading to increased membrane tension ^32^. We wondered whether the viscosity response might be impacted by Plexin-B2 function. WT and PB2 KO cells were exposed to high viscosity media for 72 hr, and confocal microscopy revealed that both showed increased F-actin, but the spread of plasma membrane beyond the cortical actin ring was farther in PB2 KO cells, consistent with less stringent attachment of plasma membrane to cortical actin (**Fig. S9C, D**).

### Molecular dynamics modeling predicts coupling of membrane involution and confined migration

Our results so far supported the importance of Plexin-B2 in orchestrating cortical actin and membrane tension during confined migration. To further understand the mechanical coupling, we performed mathematical simulations of confined migration using the molecular dynamics approach. First, we developed a coarse-grained bead model of cells with plasma membrane, actin filaments, and nuclear envelope represented by discrete bead elements connected by springs (**Fig. 5A**). The cells were then placed between two opposing virtual walls, one fixed and adhesive (simulating extracellular matrix attachment), and the other movable (simulating increasing confinement in microchannels) (**Fig. 5A**).

**Figure 5.**
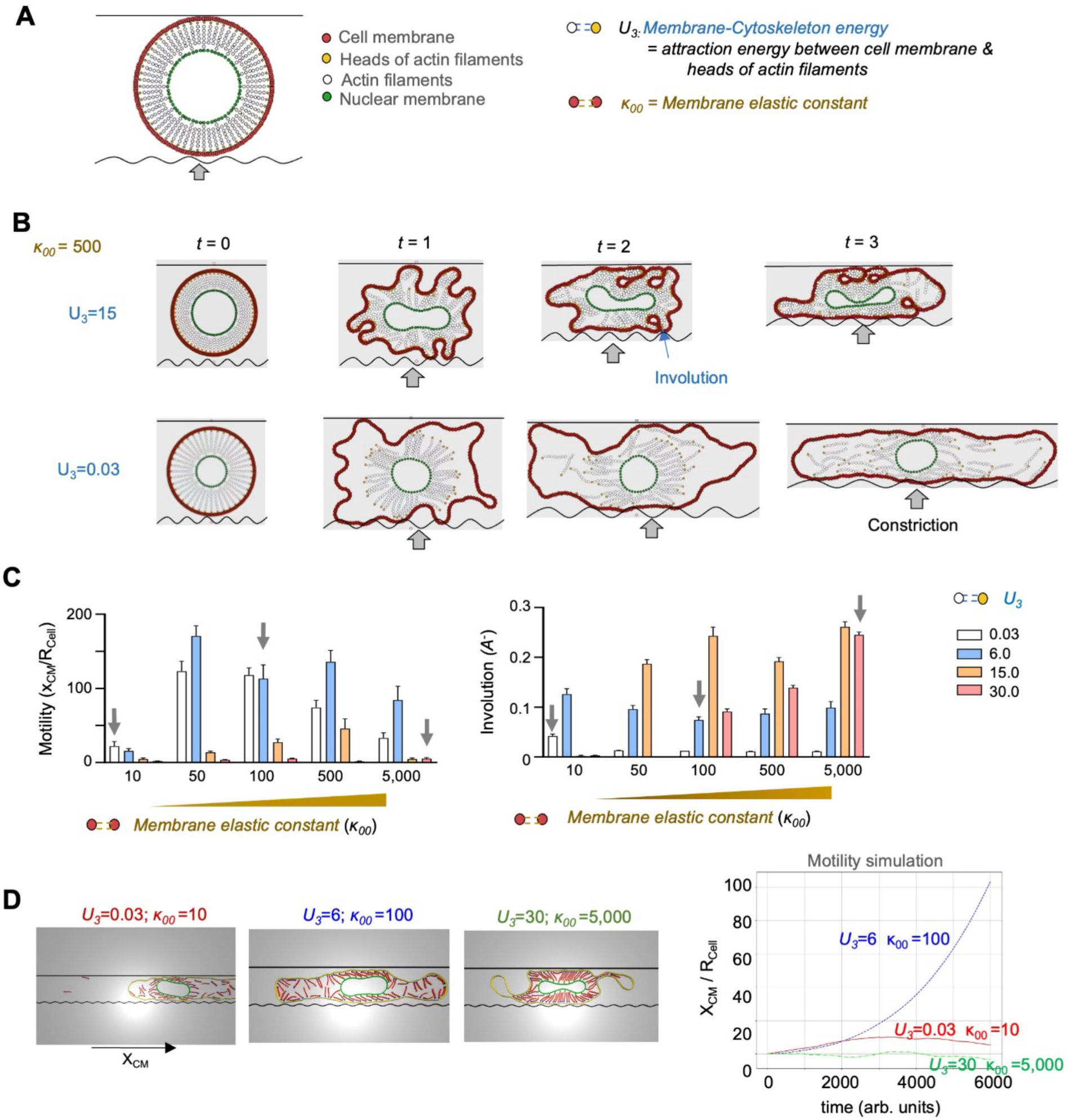
Molecular dynamics simulation illustrates the requirement of orchestrated cytoskeleton contraction and membrane tension for confined migration. (A) Cell components for molecular dynamics simulation, based on a coarse-grained bead-spring model. Two virtual walls are on opposing side of the cell, with the top wall fixed and frictionless, and the bottom wall movable with a friction factor to simulate external resistance and cell confinement. (B) Still images from simulation show cells being constricted between the virtual walls, with membrane tension set at *κ_00_*=500. At t=0, the distance between the walls is 1.2x cell diameter. At t=1 to 3, the cells are increasingly compressed. Note that for the cell with high (U_3_=15) but not low (U_3_=0.03) membrane-cytoskeleton energy condition, the plasma membrane forms involutions during constriction (arrow). See also supplemental movie S10. (C) Left, graph shows travelled distance by simulated cells (motility), measured as center of mass (CM) in the x direction (*x_CM_*) as a function of varying membrane tension *κ_00_* and membrane-cytoskeleton energy U_3_. Right, graph shows the cell negative areas (involutions; A^-^) as a function of varying *κ_00_* and U_3_ conditions. n=10 iterations per group. Data represent mean ± SEM. Arrows point to specific *κ_00_* and U_3_ conditions used for motility simulation shown in (D). (D) Left panels, still images of simulated cells migrating under constriction for three different *κ_00_* and U_3_ conditions corresponding to arrows in (C). Right panels, graphs of travelled distance by cell (motility) measured as center of mass (CM) in x direction (*x_CM_*) for three different *κ_00_* and U_3_ conditions. Note that cell motion is predicted to be minimal and lack directionality for high or low values (U_3_=30, *κ_00_*=5,000; U_3_=0.03, *κ_00_*=10), but for intermediate values (U_3_=6, *κ_00_*=100), the cell moves continuously under constriction. See also supplemental movie S11.

We first simulated confinement of cells with fixed membrane elastic constant *κ_00_* but variable membrane-cytoskeleton energy *U_3_* (approximating WT or PB2 KO conditions) (**Fig. 5B**). At *t*=0 without compression, cells in both conditions (high or low *U_3_*) assumed a round shape; but as constriction increased (*t*=1), they started to display distinct patterns of actin stretching and membrane ruffling. When constriction was further increased to simulate narrow tunnels (*t*=2 and 3), spontaneous involution of cell membrane occurred (resembling endocytic vesicles), but only in cells with high but not low membrane-cytoskeleton energy (**Fig. 5B; Movie S10**). Thus, molecular dynamics modeling predicted that cell confinement triggered membrane internalization/endocytosis requires adequate actin-membrane interaction (e.g., *U_3_=*15), resembling the WT condition.

The molecular dynamics simulations allowed us to vary both membrane-cytoskeleton energy (*U_3_*) and cell membrane elastic constant (*κ_00_*) to calculate their impact on cell motility (measured as velocity of the center of cell mass) and endocytosis (measured as plasma membrane involution or negative area A⃗^-^ (*t*)) (**Fig. 5C**). Simulations showed that confined cells displayed high motility when the membrane-cytoskeleton energy was in a medium range and cell membrane elastic constant was medium to high (κ_00_=50-5,000), indicating that cell motility is determined by balancing these forces (**Fig. 5C**). Interestingly, very high membrane-cytoskeleton energy (*U_3_*=30) did not confer higher motility, resembling PB2 OE condition (**Fig. 5C**). The involution rate of cell membrane (resembling endocytosis) followed a more direct link with intracellular forces, with highest rates of involution predicted when membrane-cytoskeleton energy *U_3_* was *≥* 15 and cell membrane elastic constant κ_00_ *≥* 50 (**Fig. 5C**).

Accordingly, mathematical simulation of cell migration over time demonstrated that intermediate values of membrane-cytoskeleton energy and cell membrane elastic constant (e.g., *U_3_*=6 and κ_00_=100) promoted sustained unidirectional confined migration over time (consistency/directional polarity) (**Fig. 5D**). Conversely, for low values (*U_3_*=0.03 and κ_00_=10), simulated cells moved for a short period in one direction but frequently changed directionality; whereas for highest values (*U_3_*=30 and κ_00_=5,000), simulated cells displayed a hypercontractile state with severely reduced motility (**Fig. 5D; Movie S11**).

### Plexin-B2 function affects membrane phospholipid composition

The lipid composition of plasma membrane may critically affect polarized migration ^7^. Anionic phospholipids –including phosphatidylinositol phosphates (PIPs) – are constituents of inner plasma membrane and crucial for signaling events ^33^. We wondered whether the reduced membrane internalization/endocytosis observed in PB2 KO cells might impact lipid composition and related signaling events.

Towards this end, we first interrogated the localization and levels of phosphatidylinositol 4,5-bisphosphate (PIP2), an essential lipid of the cell membrane and a second messenger in various signaling pathways ^34,35^. Using a fluorescent probe PH(PLCD1)-GFP, i.e., GFP fused to pleckstrin homology (PH) domain from PLCD1 that binds to PIP2, we observed that at 72 hr after transfection, the majority of GFP signal/PIP2 was internalized in WT cells, but largely retained on plasma membrane in PB2 KO cells (**Fig. 6A**). This reflected reduced endocytosis in PB2 KO cells, but may also be a sign of suppressed activity of phospholipase C (PLC), which hydrolyzes PIP2 into intracellular mediators IP3 and DAG (diacylglycerol) (**Fig. S10A**). Time course analysis revealed high PH(PLCD1)-GFP fluorescence (readout of abundant PIP2) on cell membrane of WT cells at 18 hr after transfection, which was then internalized or hydrolyzed by 24 and 48 hr; by contrast, both in PB2 KO and OE cells, the majority of PH(PLCD1)-GFP was retained in the plasma membrane, reflecting reduced internalization of PIP2/suppressed PLC activity (**Fig. S10B**).

**Figure 6.**
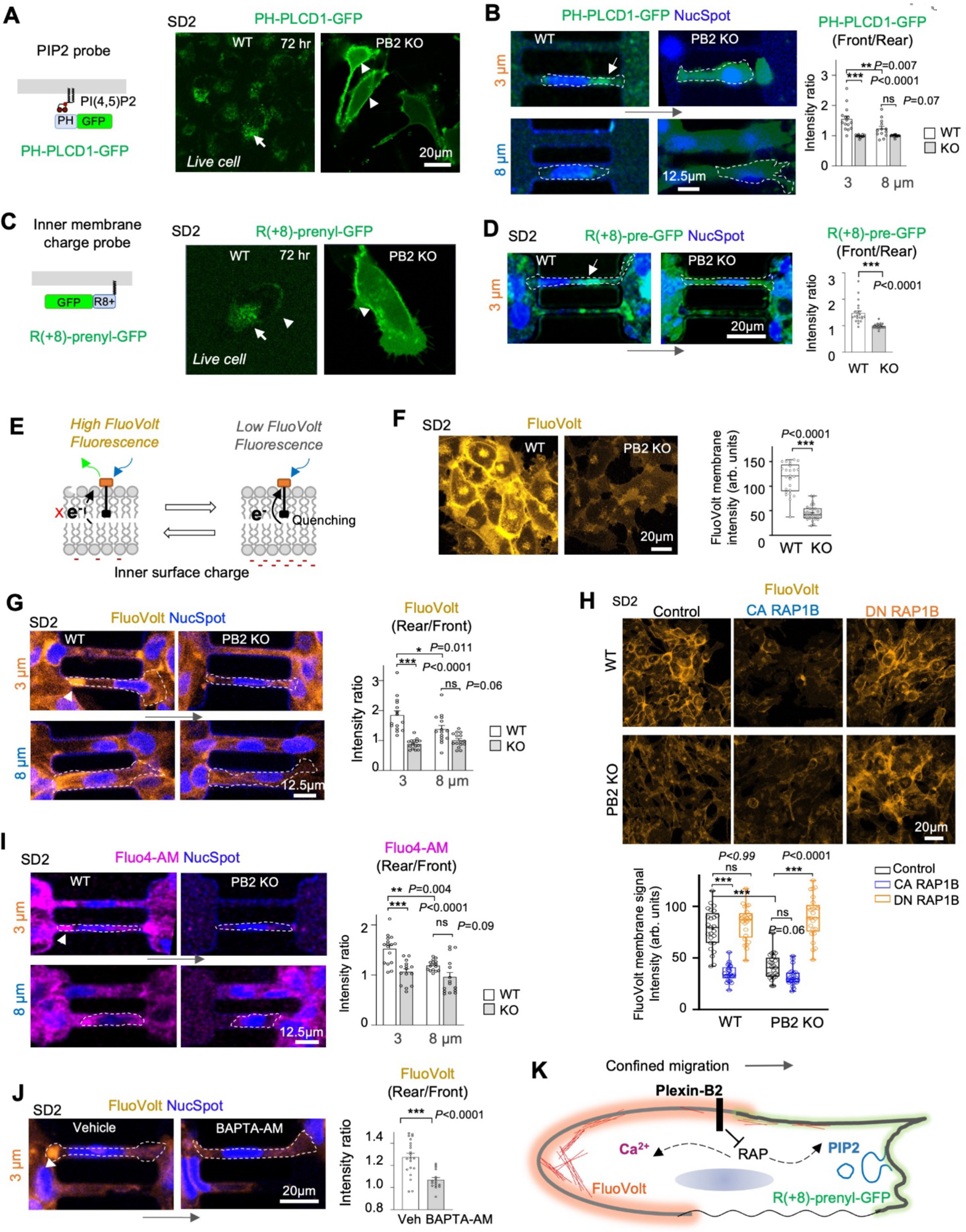
Plexin-B2 KO affects PIP2 localization and inner membrane surface charge in GSCs. (A) Left, schematic of PH(PLCδ1)-GFP PIP2 probe. Right, live-cell imaging at 72 hr post transfection reveals that the PH(PLCδ1)-GFP probes were largely internalized in WT GSCs (arrow), but retained on membrane of PB2 KO GSCs (arrowhead). (B) Left, still images of videography show accumulation of the PH(PLCδ1)-GFP probes (arrow) in front of the nucleus (NucSpot) of migrating WT SD2 GSCs in tunnels, more so in 3 than 8 µm tunnel, but not in PB2 KO cells. Dashed lines delineate cell boundary. Long arrow denotes direction of migration. Right, quantifications of the ratio of PH(PLCδ1)-GFP fluorescence intensity at front vs. rear of GSCs during passage. n=13-16 cells per condition. One-way ANOVA followed by Tukey’s multiple comparison test. Data represent mean ± SEM. (C) Left, schematic of R(+8)-pre-GFP probe for negative surface charge of inner plasma membrane. Right, live-cell imaging at 72 hr post-transfection shows internalization of the probes (arrow) in WT GSCs, in contrast to the predominant membrane localization in PB2 KO GSCs (arrowhead). (D) Left, still images of videography show accumulation of the R(+8)-pre-GFP probes (arrow) at front zone of WT GSCs when traversing the 3 µm tunnel, but not in PB2 KO cells. Right, bar graphs show the ratio of R-pre-GFP fluorescence intensity at rear vs. front of GSCs when passing through tunnels. n=22 cells for WT, n=27 cells for PB2 KO. Mann–Whitney–Wilcoxon test. Data represent mean ± SEM. (E) Diagram illustrating voltage sensitive FluoVolt membrane dye, with fluorescent intensity quenched by voltage-sensitive electron transfer from electron-rich donor mediated by “molecular wire” in plasma membrane. (F) Left, FluoVolt live-cell imaging shows reduced FluoVolt fluorescent intensity in cell membrane of Plexin-B2 KO cells, consistent with higher negative charges of inner membrane. Right, box plots of membrane FluoVolt intensity. n=25 cells for WT, n=27 cells for PB2 KO. Two-sided unpaired t-test. Data represent mean ± SEM. (G) Left, still images from videography show higher FluoVolt fluorescent signals at rear zone (arrowhead) of WT GSCs when traversing tunnels, more so in 3 than 8 µm tunnel, but not in PB2 KO cells. Migration direction is denoted by long arrow. Right, bar graphs show the ratio of FluoVolt intensity at rear vs. front during confined migration. n=15 cells per group. One-way ANOVA followed by Tukey’s multiple comparison test. Data represent mean ± SEM. (H) Live-cell images and quantifications show the effects of constitutive active (CA) RAP1B-V12 or dominant-negative (DN) RAP1B-N17 on FluoVolt intensity in WT or PB2 KO GSCs. n=25 cells per group. Kruskal–Wallis test followed by Dunn’s multiple comparisons test. (I) Left, still images capture calcium localization (Fluo4-AM fluorescence, arrowhead) at the rear of WT GSCs when traversing tunnels, more so in 3 than 8 µm tunnel, but not in PB2 KO cells. Migration direction is denoted by long arrow. Right, bar graphs showing Fluo4-AM intensity ratio at rear vs. front in GSC during passage through tunnels. n=15-16 cells. One-way ANOVA followed by Tukey’s multiple comparison test. Data represent mean ± SEM. (J) Left, still images from videography show that calcium chelator BAPTA-AM disrupted the pattern of high FluoVolt signals at the rear of WT GSCs (arrowhead) during confined migration. Right, bar graphs show FluoVolt intensity ratio at rear vs. front of GSCs when traversing tunnels. n=21 cells for WT, n=16 cells for PB2 KO. Two-sided unpaired t-test. Data represent mean ± SEM. (K) Model of Plexin-B2 signaling affecting membrane surface charge and electric field during polarized confined migration, with PIP2 enrichment at cell front and Ca^2+^ at rear zone, leading to asymmetry of FluoVolt and R(+8)-pre-GFP.

In comparison to PIP2, the membrane content of PIP3 appeared much lower in both WT and PB2 KO SD2 cells, shown by the PH(Btk)-GFP probe that binds to PIP3 ^35^ (**Fig. S10C**). This is consistent with PIP3 being a minor phospholipid constituent of the plasma membrane ^36^.

We next conducted live-cell imaging of GSCs expressing the PH(PLCD1)-GFP probe to examine PIP2 distribution during confined migration in narrow tunnels. We detected PH(PLCD1)-GFP/PIP2 accumulation at the front of migrating WT GSCs, more evident in 3 µm than in 8 µm tunnels, but not in PB2 KO cells that were stalled in the tunnels (**Fig. 6B**).

We also exposed live GSCs to fluorescently labeled annexin V in the media, which binds to phosphatidylserine (PS) this is exposed on the outer membrane ^37^. We detected internalization of annexin V-labeled endosomes at the front of WT GSCs traversing the tunnels, but not in stalled PB2 KO cells, further illustrating that membrane turnover is reduced in PB2 KO cells during confined migration (**Fig. S10D**).

### Plexin-B2 deletion alters inner membrane surface charge and local membrane electric field

PIP2 is a negatively charged phospholipid, thus contributing to the negative charge of the inner leaflet of plasma membrane ^38^. We wondered whether PIP2 accumulation in PB2 KO cells might alter inner leaflet surface charge (zeta potential) ^39^. To this end, we transfected GSCs with a plasmid expressing R(+8)-prenyl-GFP, a probe containing a series of eight positively charged arginine residues that preferentially locates to negatively charged inner membrane ^40^. We detected a distinctive pattern R(+8)-prenyl-GFP signals at cell membrane of PB2 KO cells, in contrast to WT SD2 GSCs, which harbored mostly intracellular GFP signals on endomembrane (**Fig. 6C**). Concordant with the results from the PIP2 probe, the R(+8)-prenyl-GFP probe also accumulated at the front of migrating WT cells, but not in PB2 KO cells stalled in tunnels (**Fig. 6D**). Hence, the front of confined cells contained higher levels of PIP2 and negative inner membrane charge.

As PIP2 accumulation/negative inner leaflet surface charge could impact local electric field across the plasma membrane, we labeled GSCs with FluoVolt, a voltage-sensitive fluorescent membrane dye ^41^ (**Fig. 6E**). This revealed a striking difference, with PB2 KO cells displaying lower FluoVolt signals than WT, in line with higher levels of PIP2 and negative inner leaflet surface charge in PB2 KO cells (**Fig. 6F; Fig. S11A**). In micro tunnels, migrating WT GSCs displayed higher FluoVolt signals at the rear (i.e., less negative charge), more evident in 3 µm than 8 µm tunnels, and not in stalled PB2 KO cells (**Fig. 6G; Movie S12**). The FluoVolt pattern at cell rear paralleled the F-actin pattern, but opposite that of PIP2 or membrane surface charge probe. These processes are interconnected, as endocytosis inhibitors reduced FluoVolt signals at cell rear along with compromised confined migration (**Fig. S11C; Movie S13**).

Plexins signals through intracellular GAP domain to inactivate Ras or Rap GTPases ^42^, which in turn affects a pleiotropic network of effectors controlling cytoskeletal dynamics and cell-cell adhesion ^43,44^. For further confirmation of Plexin-B2 regulation of membrane properties, we introduced constitutive active or dominant negative Rap1 mutants into GSCs (corresponding to blocked or enhanced Plexin-B2 signaling, respectively). We found that constitutively active Rap1 attenuated FluoVolt signals in WT GSCs, phenocopying PB2 KO, whereas dominant negative Rap1 reversed the PB2 KO FluoVolt phenotype (**Fig. 6H**).

We also conducted calcium imaging using Fluo-4AM, which revealed accumulation of calcium at the rear of migrating WT SD2 GSCs, more evident in the 3 µm than 8 µm tunnels, but not in stalled PB2 KO cells (**Fig 6I**). Calcium chelation by BAPTA-AM led to reduced FluoVolt signals at cell rear and compromised migration in tunnels (**Fig. 6J; S11C; Movie S13**), indicating the importance of calcium gradient for confined migration.

To control for effects that Plexin-B2 function may have on the total cell transmembrane potential, we also conducted patch clamping recordings, which showed that resting transmembrane potential and membrane conductivity were comparable between PB2 KO and WT GSCs (**Fig. S11D**). Thus, the changes of FluoVolt signals likely correspond to local changes of inner membrane surface charges.

Altogether, these results illustrate that Plexin-B2 orchestrates mechano-electric coupling, with regionalized endocytosis/PIP2/asymmetry of negative membrane surface charge at cell front, and calcium/F-actin at rear zone to propel polarized migration of GSCS through confined space (**Fig. 6K**).

### Flexible ring of Plexin-B2 extracellular domain is required for efficient confined migration

Our next question pertained to how migrating GSCs sense physical constraints and convey mechanosignals to reorganize cytoskeleton and plasma membrane. Intriguingly, a recent study unveiled that Plexin-D1 functions in endothelial cells as detector of blood flow sheer stress via its flexible ring-like extracellular domain, independent of semaphorin ligand activation ^17^. This was demonstrated by introducing a cysteines bond that prevented bending of the ring (i.e., lock mutant), thus ablating mechanosensitivity while preserving responsiveness to semaphorin. To explore whether a mechanosensitive function of Plexin-B2 might be engaged for confined migration, we introduced analogous cysteine residues into the Plexin-B2 extracellular domain to create two versions of locked ring mutants: I436C and S993C (lock1), and I436C and T1051C (lock2) (**Fig. 7A**; **Fig. S12A, B**). We first conducted control experiments in hESCs, confirming that the two lock mutants reintroduced sensitivity to semaphorin ligand but failed to rescue the Plexin-B2 KO phenotype of disrupted cellular arrangement of colonies ^19^ (**Fig. S12C-E**).

**Figure 7.**
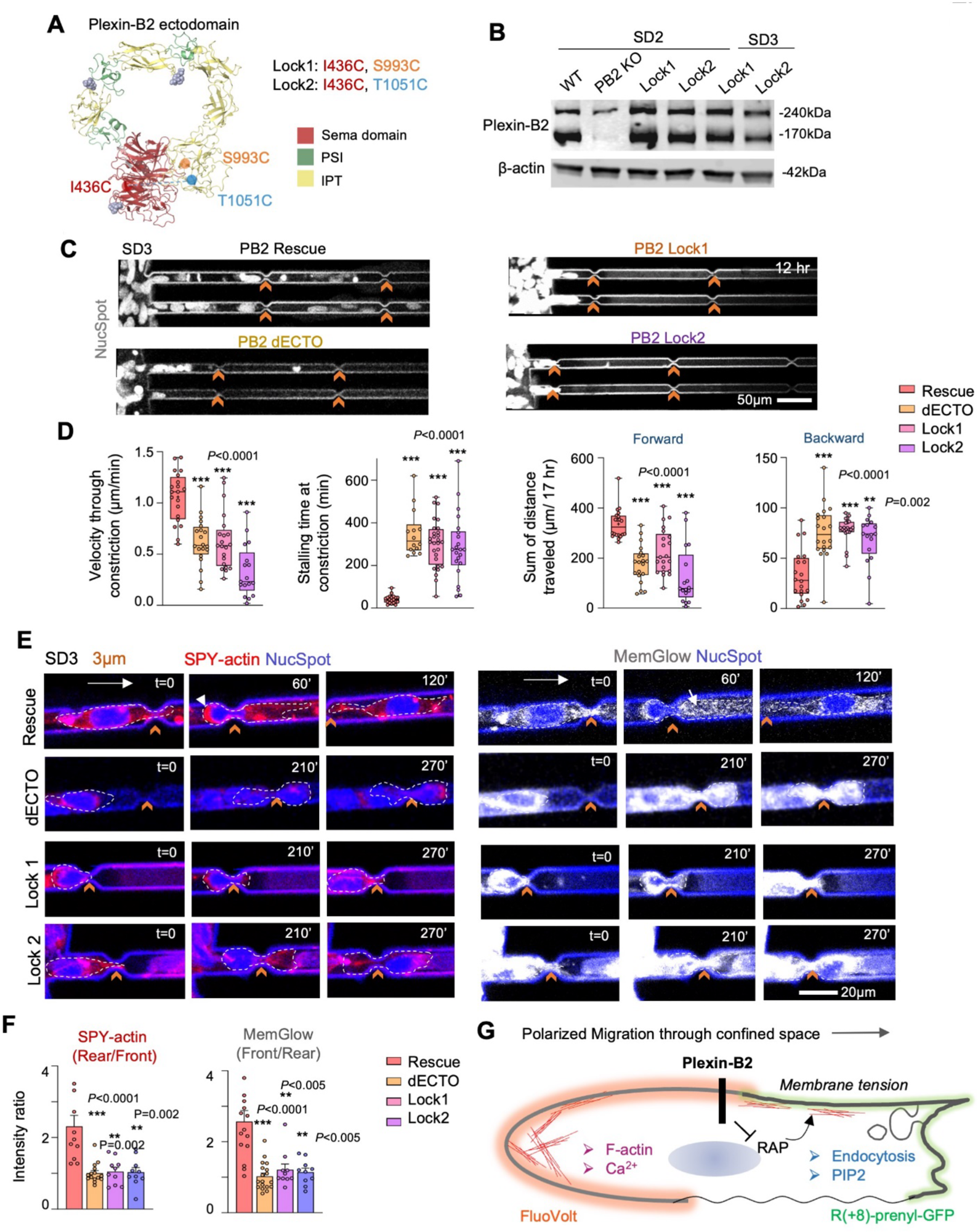
Confined migration of GSCs requires the flexible extracellular ring of Plexin-B2. (A) Structure model of the extracellular domain of human Plexin-B2 show the locations of lock1 and lock2 mutations predicted to form disulfide bridges that lock the ring structure. (B) Western blots show absence of mature Plexin-B2 (170 kDa) in PB2 KO GSC, and expression of lock mutants in PB2 KO SD2 and SD3 GSCs. β-actin serves as a loading control. (C) Still images from videography show passage of GSCs (nuclei visualized by NucSpot) through microchannels with PB2 wildtype rescue construct but not lock mutants, nor PB2 with deletion of extracellular domain (dECTO). Chevrons point to 3 µm constrictions. (D) Box plots show velocity through constrictions, stalling time at constrictions, and sum of forward and backward movements, with 25–75% quartiles, minimal and maximal values (whiskers), median (line), and mean (cross). For velocity and sum of movements: n=17-20 cells per condition. For stalling time at constriction: n=14-28 cells per condition. One-way ANOVA followed by Dunnett’s multiple comparisons test. (E) Still images from videography show F-actin assembly (SPY-actin, arrowhead) at cell rear and MemGlow^+^ endosomes (arrow) at cell front of SD3 GSCs with Plexin-B2 WT rescue but not mutant rescues when traversing 3 µm constrictions (chevrons). (F) Bar graphs showing fluorescence intensity ratio of SPY-actin and MemGlow at rear vs. front of GSCs during confined migration. n=10-18 cells per condition. Kruskal–Wallis test followed by Dunn’s multiple comparisons test. Data represent mean ± SEM. (G) Model of Plexin-B2 signaling and mechano-electrical regulation of membrane tension and membrane surface charge during polarized confined migration. Regionalized enrichment of endocytosis/PIP2 at cell front and F-actin/Ca^2+^ at rear zone lead to asymmetry of FluoVolt and R(+8)-pre-GFP membrane probes.

The lock mutants were then introduced into PB2 KO GSCs. IF staining and WB confirmed the expression of lock mutants in both SD2 and SD3 PB2 KO cells (**Fig. 7B**; **Fig. S13A**). The morphology of PB2 KO cells rescued with wildtype or lock PB2 mutants appeared comparable in 2D cultures (**Fig. S13A**). However, both lock mutants led to aberrant membrane properties, apparent as reduced endocytosis, increased membrane permeability for dextran-Alexa 488, less internalization of myr-palm-CFP membrane probes, and higher PIP2 levels in plasma membrane (**Fig. S13B-D**). Concordantly, both lock mutants resulted in lower pERM levels by WB, and reduced pERM at cell processes by IF (**Fig. S13E**), suggesting that Plexin-B2 signaling through a bendable ring to connect cortical actin to plasma membrane and cortical actin via ERM activation.

In microchannels, unlike wildtype PB2 rescue, both lock mutants and a PB2 mutant lacking the extracellular domain (dECTO) failed to rescue confined migration through 3 µm constrictions, evidenced by decreased velocity, longer stalling time, and reduced migration consistency with decreased distance of forward movement (**Fig. 7C, D; Fig. S13F**). Regionalized assembly of F-actin in rear zone and endocytosis at cell front were also disrupted (**Fig. 7E, F**).

## DISCUSSION

GBM invasion contributes to its high lethality, and understanding the cellular and molecular underpinnings is thus critical. Here, we focused on biomechanical plasticity and mechano-signaling that enable GBM cells to adapt to physical constraints to achieve polarized migration through confined space. Our results underscore a key element of membrane tension/internalization during confined migration, which is interconnected with actin contractility as well as phospholipid composition and inner membrane surface charge at regionalized zones of constricted cells. This mechano-electrical coupling and mechanosignaling are mediated through guidance receptor Plexin-B2 that orchestrates cell mechanics (F-actin assembly at rear and endocytosis at cell front), impacting phospholipid and calcium distribution, and inner membrane surface charge (**Fig. 7G**).

By utilizing 3D microdevices containing narrow constrictions and micro tunnels that mimic the physical constraints experienced by invasive GBM cells, we demonstrated that GSCs are adept at squeezing through confined space, more efficiently for 3 µm than 8 µm constrictions. Future assays can test if GSCs may perform better at constrictions wider than 3 µm (a lower limit for nucleus to pass through ^45^). Remarkably, passage through confined space further stabilized migration direction (polarity/consistency) and increased the momentum to pass through subsequent constrictions. This migratory behavior was spontaneous, driven by cell intrinsic motility, as no chemoattractant was applied. Live cell imaging captured dynamic extension of cell processes by GSCs to explore exits and rapid readjustment of membrane, cytoskeleton, and nuclear shape to squeeze through constrictions.

Our live-cell imaging also revealed robust endocytosis at cell front, which was interconnected with F-actin assembly at rear zone of constricted cells, as perturbing either process (by endocytosis inhibitors or Plexin-B2 KO to reduce actin contractility) affected the other, along with compromised confined migration. Our study supports and further extends previous report of the involvement of endocytosis in cancer cell migration ^46,47^, by highlighting regionalized endocytosis at specialized zone of confined cells (e.g., cell front). This also aligns with our molecular dynamics modeling that predicted the requirement of balanced membrane-cytoskeleton tension for optimal membrane involution and polarized migration in confined space.

The engagement of Plexin-B2-mediated endocytosis in confined migration echoes a central role of endocytosis in axon growth cone collapse triggered by semaphorins ^15,16,48–50^. Genetic studies in *C. elegans* also showed that plexin and semaphorin phenotypes could be suppressed by mutation in genes controlling vesicular transport ^51^. Our study further demonstrated membrane tension as a key feature linking Plexin-B2 and endocytosis, supported by Flipper TR imaging, AFM (gauging cell stiffness), and optical tweezers (membrane tether extension force). Our extensive interrogations of membrane properties, including dextran uptake assay, tracking of fluorogenic membrane dyes and of membrane anchored proteins (myr-palm-GFP/CFP), and the PIP2 probe (PH-PLCD1-GFP) further support a unifying model that Plexin-B2 orchestrates cortical actin dynamics and membrane tension, thereby impacting membrane internalization and permeability. Future studies using probes specific for endocytic vesicles may provide further insights.

While Plexin-B2 deletion compromised the migratory capacity of GSCs through confined space, the deformation of nuclei in narrow tunnels seemed to be unaffected by Plexin-B2 ablation. This observation suggests that proprioceptive responses of cells triggered by nuclear compression ^5^ are still intact in the absence of Plexin-B2. It is noteworthy that high viscosity-induced cortical actin contraction, which involves focal adhesions in response to mechanical stretch forces ^32^, also seemed to be independent of Plexin-B2. However, the spread of the plasma membrane beyond cortical actin network in high viscous media appeared more pronounced in Plexin-B2 KO cells, consistent with lower membrane tension and less stringent attachment of membrane to actomyosin cortex. Hence, nuclear deformation, cortical actomyosin contraction, and membrane tension could be regulated through different pathways in different physical contexts.

Our studies support the model that aside from being a receptor for class IV semaphorins, Plexin-B2 may function as a mechanosensor of compressive forces, akin to the functions of Plexin-D1 in endothelial cells and Plexin-B1 in epithelial cells ^17,18^. Plexin-B2 lock mutants with cysteine bridge introduced into the extracellular domain, thus restricting bending of the ring ^17^, failed to rescue Plexin-B2 KO phenotypes during confined migration, but was able to restore responsiveness to semaphorins. Thus, the flexibility of the ring-like structure of Plexin-B2 is key to its mechanosensitive function.

Our findings also extend the model that the negative surface charge on the inner plasma membrane acts a hub controlling cytoskeletal organization of migrating cells ^7,39,52^. The mechanisms by which Plexin-B2 influences phospholipid composition warrant further study but may involve membrane internalization or PLC activity that hydrolyzes PIP2 to IP3 and DAG, leading to calcium influx and other signaling cascades. Additionally, levels of anionic lipids like PIP2 can impact local concentration of inorganic (e.g., Ca^2+^) and organic (e.g., spermine) cations, as well as accumulation of net positively charged proteins on the inner leaflet ^53,54^, indirectly affecting cortical actomyosin assembly. The fact that calcium chelation by BAPTA reduced FluoVolt signals at rear zones of confined cells suggests an influence of calcium on electric field of local plasma membrane.

The downstream effectors of Plexin-B2 signaling during confined migration need further investigation. The conserved GAP domain of Plexin-B2 is critical for its function ^19,55^ and our FluoVolt data with Rap1B mutants suggested that Rap might be major effectors, but other Ras/Rap small GTPases such as R-Ras, M-Ras, TC21, Rin, and Rap1/2 may also be involved ^56^. Future studies are needed to test how other known drivers of migration like hypoxia, pH, and serum may impact the mechanobiology and invasion of GBM cells through confined space.

GBM is known for high intertumoral heterogeneity, and transcriptomic profiling has identified different molecular subtypes ^57^, with the mesenchymal subtype linked to aggressive infiltration and high malignant potency ^4^. Interestingly, in our assay both SD2 (mesenchymal) and SD3 (proneural) GSCs are proficient at invading microchannels, with SD3 displaying a higher velocity at constriction points than SD2. It is noteworthy that the microchannel assay does not fully equate to invasion in vivo where tumor cells need to clear extracellular matrix and interact with tumor microenvironment ^2^, nevertheless the microchannel assay provides a useful platform for dissecting mechano-coupling during confined migration.

In summary, our studies establish Plexin-B2 as a mechanoregulator of membrane tension that facilitates confined migration of GBM cells. Understanding how membrane mechanics and mechano-electrical coupling enable GBM cells to adapt to physical constraints and rapidly adjust migratory behavior may help open new therapeutic avenues to curb GBM invasion.

## METHODS

### Human GBM cell lines

De-identified human GBM stem cell (GSC) lines SD2 and SD3 had been established from resected tumor tissue of GBM patients at the University of California, San Diego, and have been characterized by their transcriptomes as mesenchymal and proneural subtype, respectively ^6^. GSCs were propagated in neural stem cell media (Neurocult NS-A proliferation kit (human), Stemcell Technologies), supplemented with bFGF (10 ng/ml; Peprotech), EGF (20 ng/ml; Peprotech), heparin (0.0002%; Stemcell Technologies), and penicillin–streptomycin (1:100; Gibco). Cells were propagated in adherent conditions on culture dishes coated with laminin (10 µg/ml in PBS; 1 hour at 37 °C; Gibco). Passaging of GSCs was performed by dissociating cells with Accutase (BD Biosciences).

### Neural differentiation

Human neural progenitor cells (hNPCs) were generated by a single-cell monolayer protocol using STEMDiff Neural Induction Medium (StemCell Technologies) as described previously ^19^. Briefly, hESCs were plated at a density of 2.5 × 10^5^ cells/cm^2^ in STEMDiff medium, and on day 6 cells were replated at the high confluence. After day 11, cells were plated on glass coverslips coated with Matrigel for analysis by immunocytochemistry to confirm NPC identity and were then used for subsequent experiments using microchannels devices.

### Microchannel migration device

To mimic the passage of GBM cells through narrow pores, microchannel devices consisting of polydimethylsiloxane (PDMS) polymer structures on 35 mm glass bottom Petri dishes with parallel rows of microchannels of 10 µm height, 12 µm width, and constrictions of 3 or 8 µm were used (4D Cell, #MC011).

To simulate the passage of GBM cells through narrow tunnels, we created microdevices containing central chambers with outflow tunnels of 3 or 8 μm width and 50 µm length. This device allows cells to choose different exit tunnels from the chambers. These devices were designed using CAD software to create a photolithographic chrome mask for groove and cell culture compartment regions. Photolithography was then used to print mask features onto a silicon wafer, coated with negative SU-8 photoresist. The final devices were fabricated in a replica molding process by casting a PDMS prepolymer mixture against the positive relief master mold to obtain a negative device replica. Well openings were punched into devices, which were then bonded to the glass bottom of a cell culture dish using corona discharge treatment.

Before seeding of cells, the devices were immersed in 70% ethanol and left for 5 min inside a benchtop vacuum desiccator (Southern Labware) to remove air bubbles. After washes with PBS, the microchannels were coated with laminin (100 µg/ml in PBS; Invitrogen) for 1 hour at 37 °C. All wash and coating steps included 5 min placement inside a vacuum desiccator.

To initiate the microchannel culture, 3×10^4^ - 10^5^ GSCs cells were seeded into the entry ports. After incubation for 4 - 24 hours, cells were labeled by adding the dyes MemGlow 488 (Cytoskeleton; 1:200), SPY555-Actin (Cytoskeleton; 1:1,000), and NucSpot Live 650 (Biotium; 1:1,000). Migrating GBM cells in the microchannels were imaged every 5 min on a LSM 780 (Zeiss) confocal microscope (heated stage, 37 °C, with 5% CO_2_) for up to 17 hr. Image analysis was performed with ImageJ (MTrackJ plugin) to determine values as described below.

For the devices containing microchannels with constrictions, velocity of cells through constrictions was calculated by tracking cells from the time the front of the cell reached the constriction (position 1) until the rear of the cell completely exited the constriction (position 2). The time and distance between the two positions were used to calculate velocity. Stalling time was calculated as the time span a migrating cell would remain stalled in front of a constriction before entering the constriction.

To calculate the ratio of intensities of different fluorescent labels, including SPY-actin, Fluo-4, FluoVolt, and PH-PLCD1-GFP, annexin V, and MemGlow, the different front and rear fluorescence intensities of a migrating cell were measured when the nucleus was in the middle of a tunnel or constriction. The mean intensities were measured within each compartment (using the nucleus as boundary between the front or rear compartments).

### Treatment with endocytosis inhibitors

Treatment with small molecule endocytosis inhibitors for live-cell imaging experiments was carried out on cells plated into the microchannel devices. Cells were incubated for 10 min with Pitstop 2 (30 µM; Abcam)^24^, 30 min with Dynasore (80µM; Sigma)^58^, or 90 min with 5-(N-ethyl-N-isopropyl)-amiloride (EIPA) (50µM; Cayman Chemical)^59^. After drug or vehicle treatment, the GBM cells were washed with PBS and stained with SPY555-Actin, NucSpot, and FluoVolt or MemGlow as described above and imaged immediately on a LSM 780 confocal microscope (Zeiss).

### Scratch assay

GBM cells were seeded in six-well plates at a density of 10^6^ cells per well. After one day of culture, cells were treated with endocytosis inhibitors (Pitstop 2, Dynasore, and EIPA) as described above. The cells were washed with PBS and the media was replenished, followed by a longitudinal scratch made at the center of the dish with a 200-µl size micropipette tip (t = 0 hr), and wound closure speed was determined by micrographs of identical areas at t = 24 hr, 48 hr, and 72 hr after scratch.

### Immunocytochemistry

For immunocytochemistry (ICC) of cells in culture, cells were fixed with 3.7% formaldehyde in PBS at room temperature for 10 min, followed by three washes with PBS, and incubation with blocking buffer (5% donkey serum and 0.3% Triton X-100 in PBS) for 1 hr. Primary antibodies were added in dilution buffer (PBS with 1% BSA and 0.3% Triton X-100) at 4 °C overnight. Secondary antibodies were incubated in dilution buffer at room temperature for 1 hr together with nuclear stain (DAPI, 1:1000; Thermo Fisher). For most preparations, the cells were also counterstained for F-actin with Alexa-594 phalloidin (Thermo Fisher).

### Western blotting

For Western blotting (WB), cells were lysed with RIPA buffer (Sigma) containing protease and phosphatase inhibitors. For phospho-ERM WB, cells were lysed using Cell Lysis Buffer (Cell Signaling, 9803). Protein concentrations were determined using a BCA assay (Thermo Scientific). Proteins were resolved by SDS-PAGE on 4–12% polyacrylamide NuPAGE gels (Invitrogen) and transferred onto nitrocellulose or PVDF membranes (Li-Cor Biosciences) with the XCell transfer system (Invitrogen). Membranes were incubated at 4 °C overnight with primary antibodies and then for 1 hr with secondary donkey antibodies coupled to IRDye 680 or 800 (Li-Cor). The fluorescent bands were detected with an Odyssey infrared imaging system (Li-Cor Biosciences).

### Antibodies

Primary antibodies for ICC:

anti-Plexin-B2, extracellular domain, R&D systems AF5329 (sheep), dilution 1:300;

anti-Ezrin, Sigma SAB4200806 (mouse), dilution 1:1,000;

anti-phospho-ERM (Ezrin (Thr567)/Radixin (Thr564)/Moesin (Thr558)) (48G2), Cell Signaling 3726S (rabbit), dilution 1:1,000;

anti-Ki67, Abcam ab15580 (rabbit), dilution 1:200.

Secondary antibodies for ICC:

Alexa Fluor 488, 594, or 647-conjugated donkey anti-sheep, -rabbit, or -mouse IgG (Jackson ImmunoResearch Laboratories, dilution 1:300).

Primary antibodies for WB:

anti-Plexin-B2, extracellular domain, R&D systems AF5329 (sheep), dilution 1:500;

anti-Ezrin, Sigma SAB4200806 (mouse), dilution 1:1,000;

anti-phospho-ERM (Ezrin (Thr567)/Radixin (Thr564)/Moesin (Thr558)), Cell Signaling 3141S (rabbit), dilution 1:500;

anti-β-actin, Sigma A1978 (mouse), dilution 1:10,000.

Secondary antibodies for WB:

IRDye 800CW anti-mouse (donkey, Li-Cor Biosciences 926-32212,1:10,000), IRDye 680LT anti-goat (donkey, Li-Cor Biosciences 926-68024, 1:10,000), and IRDye 680RD anti-rabbit (donkey, Li-Cor Biosciences 926-68073, 1:10,000).

### CRISPR KO of *PLXNB2*

The deletion of Plexin-B2 in human cell lines by lentiviral CRISPR-Cas9 vectors has been described previously ^6,19^. Briefly, GSCs were stably transduced with lentivirus expressing Cas9 and a short guide RNA against the second coding exon of *PLXNB2* or a guide RNA against the EGFP coding sequence (used as control). The plasmids for the lenti-CRISPR vectors have been deposited at Addgene (#86152 and #86153, respectively).

### Lentiviral Plexin-B2 cDNA vectors

For overexpression and rescue experiments, human *PLXNB2* cDNA lentiviral vectors described previously ^6,19^ were used to transduce GSCs. The plasmids for lenti-*PLXNB2* expression have been deposited at Addgene (pLV-PLXNB2: #86237 and pLV-PLXNB2-dECTO).

The locations of the Plexin-B2 lock mutations were determined by structural prediction of Plexin-B2 as described below. The mutations I436C & S993C (“Lock1”) and I436C & T1051C (“Lock2”) were introduced into a CRISPR-resistant *PLXNB2* cDNA plasmid by site directed mutagenesis (Thermo). The mutant cDNAs were transferred by Gateway reaction into pLENTI PGK Neo DEST (Addgene #19067; deposited by Eric Campeau & Paul Kaufman). The lock mutant plasmids have been deposited at Addgene (lock1: #182879 and lock2: #182880). The Plexin-B2 lock mutant expressing lentiviruses were transduced into GBM cells carrying CRISPR/Cas9 Plexin-B2 KO and were selected with 200 µg/ml G418 (Gibco) for 7 days before confirmation of expression by WB and ICC.

### Prediction of Plexin-B2 structure for locked ring mutations

We first predicted the 3D structure of human Plexin-B2 by mapping it onto multiple templates, using methods as described previously ^12^. To model the extracellular part of HsPLXB2, we used the PDB protein structure 5L56 as template, and to model the intracellular part, we used the PDB structures 5E6P, 3IG3, and 3RYT as templates. We identified amino acid positions in the N-terminal Sema domain and the juxta-membrane IPT5 domain where newly introduced cysteines are prone to form a disulfide to lock the ring. The mutations in I436C, S993C, and T1051C were obtained using the mutagenesis tool of Pymol and optimized through Swiss PDB Viewer. To determine these mutations, we analyzed the consensus results of the following predictors: (i) Disulfide by Design (cptweb.cpt.wayne.edu/DbD2/index.php), (ii) CYSPRED (gpcr.biocomp.unibo.it/cgi/predictors/cyspred/pred_cyspredcgi.cgi), (iii) Protein Interactions Calculator (pic.mbu.iisc.ernet.in/job.html), and (iv) PDB2PQR and PROPKA programs (server.poissonboltzmann.org/pdb2pqr).

### Doxycycline-inducible Plexin-B2 knockdown or overexpression

Temporally controlled knockdown of Plexin-B2 was achieved with TET-ON lentiviral vectors expressing doxycycline (Dox)-inducible shRNA targeting *PLXNB2* (pLKO-Tet-On-*PLXNB2*–shRNA2; deposited as Addgene #98400). Puromycin selection at 1 µg/ml was used to establish transduced cell lines.

The lentiviral vector for Dox-inducible Plexin-B2 overexpression was generated by inserting human *PLXNB2* cDNA into a Dox controlled expression vector (pLenti-CMVtight-PLXNB2 iOE; deposited as Addgene #176849) ^19^. Target cells were coinfected with a lentivirus expressing Tet-On 3G transactivator protein with hygromycin resistance (VectorBuilder). Stable *PLXNB2* Tet-On hESC lines were established by successive puromycin (1 μg/ml) and hygromycin (200 μg/ml) selection steps. The expression of *PLXNB2* was induced by addition of 1 μg/ml Dox (MP Biomedicals) to the culture medium.

### GBM cell elasticity measured by atomic force microscopy

Atomic force microscopy (AFM) measurements were conducted using an Asylum MFP-3D-BIO device (Asylum Research), coupled to an Olympus IX-80 inverted microscope, similar to methods described previously ^19^. Briefly, for AFM experiments, 10^5^ GBM cells were plated on 60 mm laminin-coated tissue culture dishes. The AFM setup included a gold-coated silicon nitride probe with triangular shaped body, blunted pyramidal tip (39° half-angle), and nominal spring constant of 0.09 N/m (TR400PB, Asylum Research). A 3×3 indentation array was conducted over a 5 µm square region of each cell body, avoiding the nucleus and cell edge. The AFM was set to perform indentations at a rate of 1 Hz and trigger at 20 nm of cantilever deflection to ensure consistency across measurements. Each indentation was fitted using a Hertzian model (assuming cell Poisson’s ratio of 0.45) to calculate the modulus of elasticity. Poorly fitted curves were removed from downstream analysis.

### Measurement of membrane tension with Flipper-TR dye

Membrane tension measurements with the dye Flipper-TR (Spirochrome SC020) ^29^ were performed by labelling GSC lines with Flipper-TR (1:1,000) in neural stem cell media. After incubation at 37 °C for 15 min, cells were washed with PBS and once with fresh media, and cells were imaged live at a Leica TCS SP8 STED 3X super-resolution microscope on a heated and CO_2_ and humidity-controlled stage. Z-stack images (1 µm total height, 0.1 µm intervals) were acquired from cell membrane regions using a 93x Plan-Apochromat 1.3 NA glycerin immersion objective. Excitation was performed using a pulsed 488 nm laser operating at 20 MHz with emission collected through bandpass 565/610 nm filter. Lifetimes of Flipper-TR were extracted from fluorescence lifetime microscopy (FLIM) images and fitted to a dual exponential model using FLIM Wizard LAS AF software (Leica). The longest lifetime with the higher fit amplitude was used to quantify membrane tension (lifetime (τ2) at the Leica LAS software).

### Measurement of membrane tension with optical tweezers

The measurement of membrane tension with optical tweezers was performed using a C-Trap device (Lumicks). To measure the mechanical properties of the cell membrane, carboxyl polystyrene particles (1.5 – 1.9 µm diameter, Spherotec) were coated with the lectin concanavalin A (eBioscience) and added to the culture medium. A laser beam (10 W, 1,064 nm) was focused through a series of Keplerian beam expanders and a high numerical aperture objective (×63/1.45, oil immersion, Nikon) to trap a particle, which was then moved towards a cell membrane at a speed of 1 µm/s. After momentary contact between bead and plasma membrane, beads were moved away again to extend tethers from cells. The displacement of the bead from the center of the optical trap was recorded by a position-sensitive detector to calculate the tension force. Cell membrane stiffness was determined as the stable force value that was obtained during tether extrusion. Data collection and post-processing were performed using Lumicks Bluelake software and Python.

Optical tweezers measurements after treatment with drugs targeting the cytoskeleton were carried out directly in live GSCs after 3 hr of addition of blebbistatin (10 μM), latrunculin (5 μM), or vehicle. After the experiment, GSCs were fixed and stained with phalloidin for F-actin and DAPI for nucleus and imaged using LSM 710 confocal microscope (Zeiss).

### Whole-cell patch-clamp recordings

Cells were plated onto poly-D-lysine (100 mg/ml; SigmaAldrich) coated 12-mm glass coverslips in 24-well plates. Whole-cell patch-clamp recordings were made as described previously ^60^. Borosilicate glass electrodes (Warner Instruments) of 3-6 MΩ were filled with an intracellular solution containing 130 mM KCl, 20 mM NaCl, 5 mM EGTA, 5.46 mM MgCl_2_, 2.56 mM K_2_ATP, 0.3 mM Li_2_GTP and 10 mM HEPES (pH 7.4, ∼ 320 mOsm). The extracellular solution contained 5 mM KCl, 155 mM NaCl, 0.5 mM CaCl_2_, 2 mM MgCl_2_, and 10 mM HEPES (pH 7.4, ∼325 mOsm). For voltage clamp recordings, currents were elicited at 0.5 Hz with voltage steps to -100 mV from a holding potential of -70 mV, then repeated to the next voltage steps until + 80 mV (20 mV difference in each step). For current clamp recordings, currents were held at 0 A, and the changes in membrane potential (Vm) were measured.

### Fluorescent protein localization assays

Plasmids for expression of lipid anchored fluorescent proteins were obtained from the Addgene repository: MyrPalm-CFP (Addgene #14867) and MyrPalm-GFP (#21037), as well as plasmids for expression of a PI(4,5)P2 binding fluorescent protein: PH-PLCD1-GFP (#51407) and a PIP3 binding protein: PH-Btk-GFP (#51463). Also obtained from Addgene was a plasmid expressing a surface charge probe containing a series of arginine residues linked to a prenylation signal fused with GFP R(+8)-prenyl-GFP (#17274).

For plasmid transfection, a cell suspension of GBM cells was electroporated using a Neon transfection system (Invitrogen) with the following parameters: 2 × 10^5^ cells, 15 µg of plasmid DNA, 1 pulse for 30 ms at 1,300 V. Cells were seeded after electroporation on laminin-coated four-chamber glass-bottom dishes (Cellvis) at a density of 5×10^4^ cells per chamber and live imaged using a LSM 710 confocal microscope (Zeiss).

### Dextran-Alexa 488 endocytosis assay

To visualize the endocytic activity of GBM cells, cells were incubated for 40 min with 5 mg/ml dextran Alexa488, MW 10,000 (Invitrogen D22910). After a wash with PBS, live cells were imaged at a confocal microscope. Dextran was excited at 488 nm and emitted fluorescence was recorded with an LSM 710 confocal microscope (Zeiss).

Low temperature effect on membrane permeability was assessed by keeping cells at 4 °C for 10 min, followed by dextran treatment also at 4 °C for 10 min, and then immediate imaging at a LSM 710 confocal microscope (Zeiss).

To visualize the effects of cytoskeletal drug treatment on dextran Alexa488 GSC permeability, cells were incubated for 1 hr with blebbistatin (20 μM). After wash with PBS, GSCs were incubated with dextran Alexa488 as described above, stained with nuclear dye Hoechst 33342 (1:2,000; Thermo 62249) and imaged using LSM 710 confocal microscope (Zeiss).

### Staining of cells for cytoskeleton, lysosomes, and membrane components

To visualize filamentous actin in live cells, we stained cells with SPY555-actin (1:1,000; Cytoskeleton CY-SC202) at 37 °C for 1 hr. To visualize lysosomes, cultured cells were stained with LysoTracker Red DND-99 (Invitrogen L7528; 75 nM) at 37 °C for 10 min. To assess cell viability, cultured cells were stained with membrane permeable Calcein Violet 450 AM (Thermo; 5 µM) at 37 °C for 20 min.

### Membrane dynamics and endocytosis assessment with MemGlow, NR12A, and Flipper-TR

For the assessment of membrane dynamics and endocytosis, GBM stem cells were stained with MemGlow or NR12A for 20 min at 37 °C then imaged on a LSM 710 or 780 confocal microscope (Zeiss). For temporal analysis of endocytosis, GSCs were stained with Flipper-TR for 20 min at 37 °C and cells were analyzed at 2 -5 hr time points using Leica TCS SP8 STED 3X super-resolution microscope at 37 °C and 5% CO_2_. No wash step was used for these experiments.

### Three-dimensional surface rendering of GSCs

Z-stack images (20 µm total height, 0.21 µm intervals) were acquired from live-cell membranes stained with Nile red (NR12A) dye using a Plan-Neofluar 40x/1.30 oil immersion objective on a LSM 710 confocal microscope (Zeiss). Three-dimensional surface rendering was created in surpass 3D mode from CZI files (Zeiss software) using Imaris v10.0 (Bitplane).

### FluoVolt live staining

To assess membrane potential of cells in 2D cell culture, FluoVolt (Thermo, 1:1,000) membrane dye was diluted together with Powerload supplement (1:100) in neural stem cell media according to manufacturer’s instructions. Cells were stained for 30 min at 37 °C in a cell culture incubator, washed twice with PBS, and then imaged 15 min later with 488 nm excitation with an LSM 710 or 780 inverted confocal microscope (Zeiss).

To assess membrane potential of cells migrating in microchannels, cells were seeded in laminin-coated microchannel devices at a density of 3 × 10^4^ cells per chamber. After one day, cells were incubated with FluoVolt dye diluted together with Powerload supplement as described above. Additionally, NucSpot Live 650 (1:1,000; Biotium) dye was used to label nuclei. Cells were imaged on a Zeiss LSM 780 inverted confocal microscope.

### Annexin V live phosphatidylserine staining

To detect the negatively charged lipid phosphatidylserine (PS), cells were seeded in laminin-coated microchannel devices at a density of 3 × 10^4^ cells per chamber. After one day of culture, cells were incubated with fluorescent PS binding protein annexin V-Alexa Flour 488 (1:10; Thermo A13201) in annexin-binding buffer for 15 min at room temperature. Cells were then washed with annexin-binding buffer and additionally incubated with NucSpot Live 650 (1:1,000; Biotium) for 1 hr at 37 °C. Cells were imaged at 488 nm excitation and emitted fluorescence was recorded with an LSM 780 inverted confocal microscope (Zeiss).

### Fluo-4 live staining

To visualize Ca^2+^ signals, cells were seeded in laminin-coated microchannel devices at a density of 3x 10^4^ cells per chamber. After one day of culture, cells were incubated with NucSpot Live 650 (1:1,000; Biotium), the fluorescent Ca^2+^ indicator Fluo-4-AM (1:500; Invitrogen), and 20% pluronic acid F-127 (1:500; Thermo P3000MP) for 1 hr at 37 °C. Cells were kept in medium and imaging was performed at 488 nm excitation for Fluo-4-AM and 647 nm excitation for NucSpot and emitted fluorescence was recorded with an LSM 780 inverted confocal microscope (Zeiss).

### Viscous media assay

Viscous media was prepared by adding methylcellulose 65 kDa (Sigma) at a concentration of 0.6% (92 μM) to neural stem cell media to obtain a viscosity of 8 centipoise (cP). Notably, this is in the maximum range of physiological viscosity ^61^. To assess the effects of viscosity on the cytoskeleton and membrane dynamics of GBM cells, the cells were plated in regular neural stem cell media for 24 hr and exposed to 8 cP viscous media for 72 hr. The control group was kept in regular neural stem cell media. The cells were labeled with the dyes SPY555-Actin (Cytoskeleton; 1:1,000) and NucSpot Live 650 (Biotium; 1:1,000) for 1 hr. The media was replaced with fresh media containing MemGlow 488 (Cytoskeleton; 1:200) and cells were incubated for 20 min at 4 °C. GBM cells were then imaged on a LSM 710 (Zeiss) confocal microscope. The areas of plasma membrane (MemGlow) and of actin filaments (SPYactin) were quantified using ImageJ, and the ratio of membrane vs. actin areas was reported as membrane spreading.

Relative SPY555-actin and MemGlow intensities were measured using the corrected total cell fluorescence quantification (CTCF) method with ImageJ. Briefly, cells were manually selected on micrographs, and CTCF for SPY555-actin and MemGlow were calculated for each cell as (integrated density of fluorescence signal) − (area of cell × mean fluorescence background reading).

### Treatment with oleoyl-L-α-lysophosphatidic acid

Treatment with oleoyl-L-α-lysophosphatidic acid (LPA; 5 µM; Sigma, L7260) was carried out by addition to media of live cells for 5 min. After treatment, GSCs were washed with PBS and fixed for ICC or lysed in RIPA buffer containing protease and phosphatase inhibitors for subsequent Western blotting analysis.

### Molecular dynamics simulations

#### Cell model

Coarse-grained molecular dynamics modelling was performed using a cell model composed of beads representing the cell membrane, the nuclear membrane, actin filaments, and heads of actin filaments, as described previously ^19^. The nuclear membrane is connected to the cell membrane through actin filaments. Connections between the beads are modelled by springs of elastic constant κ. The total number of beads per cell in the model was n=700.

#### Potentials

Each bead was associated with potentials, which produce an interaction force with neighbor beads. Thus, these potentials are responsible for generating the dynamics of the system. As we are differentiating cell membrane (particle 0), actin (particle 1), and heads of actin filaments (particle 3), the elastic constants associated with each particle may be different. Thus, κ_00_ is the elastic constant that connects the membrane (particles 0 with 0), κ_30_ is the constant that connects particle 3 with particle 0. The model allows determining the position vector 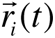 of each bead *i* at time *t* in relation to a coordinate system. With the position vector it is possible to find the position vector of the center of mass in the x direction, *x*_CM_(*t*). The model also allows, through the position vectors, to determine the area vector 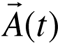 of the cell at each time point. The negative areas correlated with membrane involution.

To compute the forces of interactions between a given particle *i* and its neighbors we chose some common potentials in the literature in the study of polymers. The potentials used in the model are Lennard-Jones Potentials (LJ), Weeks-Chandler-Anderson Potentials (WCA), Finite Extensible Nonlinear Elastic (FENE), and Bend Potential (BP). The potential BP allows the twisting of membranes and actin filaments. Thus, the total potential to which each particle *i* is subjected is given by:

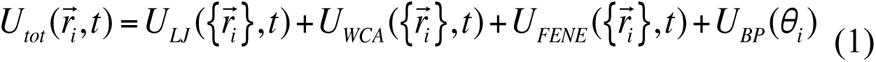

where θ*_i_* is the angle formed between 3 particles, the middle particle being particle *i* and 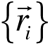 represents the set of position vectors of particle *i* with all other particles. With the expression of the potential, it is possible to obtain the force that is subject to particle *i* is given by

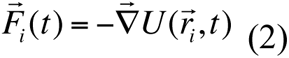

Constrictions simulations: To perform constriction simulations, we added two walls to the model, one mobile and the other fixed. The fixed wall is at the top and there is no friction. The bottom wall is movable and has friction. The potential associated with friction was modeled by a cosine function in the x direction, that is,

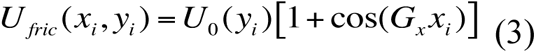

where *G_x_* = 2*π* / *a* is the lattice parameter and *a* being the distance between two beads that make up the bottom wall. The potential *U_0_(y_i_)* is of the type:

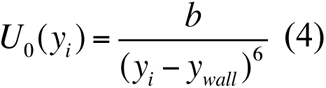

where *y_i_* is the vertical position of a given particle *i*, *y_wall_* is the vertical distance from the wall to that particle and *b* is a constant. This potential (equation 3) must be added to the total potential of equation 1. The frictional force generated by this potential produces an action-reaction pair that allows, under certain conditions, the cell to move between the walls. Initially *(t = 0*) the cell has a circular shape with radius R_C_ and the walls are farther apart. The system evolves and close to time *t =2* (*250 arb. units*) the lower wall stops moving, remaining at a distance *d < R_C_* of the top wall. At this instant, the cell is constricted between the upper and lower walls.

For the constriction simulation, the variables *N, ϒ* (representing the ratio between nucleus and cell radius)*, R_C_, U_fric_* and *d* were fixed. The variables of interest were the elastic constant of the membrane (κ_00_) representing the membrane tension and the potential for the head of the actin filament to bind to the cell membrane beads, which were defined as *U_3_*.

Thus, we used this model to analyze the movement of the cell’s center of mass (CM) and area (*A*) as a function of time for a given set of parameters of cell membrane elastic constant κ_00_, and the potential at the head of the actin filament that binds to the cell membrane (*U_3_*).

### Statistics and Reproducibility

Data are presented as mean ±SEM unless indicated otherwise. One-way analysis of variance (ANOVA) with Tukey’s or Dunnett’s post-hoc test or Kruskal-Wallis test with Dunn’s multiple comparisons test were performed for data with more than two groups. Unpaired *t*-test (two sided) or Mann-Whitney test was applied when comparing data from two groups. For studies with repeated measures (RM), two-way ANOVA RM and Bonferroni post-hoc test was performed. Statistical analyses were performed with GraphPad Prism 9 statistical software, using the style setting NEJM (New England Journal of Medicine) for reporting of *P* values. Statistical significance was considered as *P*<0.05 (*); *P*<0.01 (**); *P*<0.001 (***). Sample sizes and statistical details are indicated in figure legends. All cell culture experiments presented in the manuscript were repeated at least three times independently with similar results.

## DATA AVAILABILITY

All data that has been generated for this study are available from the corresponding author R.H.F. upon reasonable request.

## CODE AVAILABILITY

Molecular dynamics (MD) codes have been deposited at Github (github.com/diasrodri/SimCellMD-1) and Zenodo (https://zenodo.org/doi/10.5281/zenodo.4977916).

## Supporting information

Movie S1 SD2 3 vs 8 um constriction (related to Fig. 1)

Movie S2 SD2 vs SD3 migration (related to Fig. 1)

Movie S3 NPC vs GSC migration (related to Fig. 1)

Movie S4 Endocytosis inhibitors (related to Fig. 1)

Movie S5 NR12A membrane projection (related to Fig. 2)

Movie S6 SD2 WT vs PB2 KO (related to Fig. S8)

Movie S7 SD3 WT vs PB2 KO (related to Fig. 4)

Movie S8 tether microchannel (related to Fig. 4

Movie S9 SD3 Chamber WT vs PB2 KO (related to Fig. 4)

Movie S10 Molecular dynamics Endocytosis (related to Fig. 5)

Movie S11 Molecular dynamics Movement (related to Fig. 5)

Movie S12 SD2 Chamber FluoVolt WT vs PB2 KO (rel to Fig. 6)

Movie S13 SD2 Chamber BAPTA treatment (rel to Fig. 6)

## ACKNOWLEDGEMENTS

We thank Yvonne Jones and Vitul Jain, University of Oxford, for advice on Plexin structure predictions. This work was supported by a National Institutes of Health K01 career development award to C.J.A (NS127948) and research awards to R.H.F (NS092735, NS125700) and H.Z. (NS107462, NS134159). Additional support was provided by New York State Department of Health awards to H.Z. (C38330GG and C39068GG) and CAPES and CNPq funding to R.A.D, J.P.M, and P.V.Z.C.. Fellowship support was provided by FAPEMIG and UFJF to J.S.L. and P.F..

## AUTHOR CONTRIBUTIONS

C.J.A., R.H.F., and H.Z. designed the overall study, analyzed, and compiled data, and wrote the manuscript. C.J.A. and T.H. conducted experiments and performed data analyses. R.J.W. and K.C. performed AFM measurements and analyzed the data. H.N. and P.S. performed and analyzed the electrophysiological measurements. M.J.T. performed the OT experiment. J.S.L., P.F.C.F, N.P.D.N., and P.V.Z.C. performed the MD simulations for predicting PB2 LR mutations. R.A.D. and J.P.M. performed mathematical modeling. All authors discussed manuscript preparation.

## References

1. Cuddapah, V.A., Robel, S., Watkins, S., and Sontheimer, H. (2014). A neurocentric perspective on glioma invasion. Nat Rev Neurosci 15, 455–465. 10.1038/nrn3765.

2. Mair, D.B., Ames, H.M., and Li, R. (2018). Mechanisms of invasion and motility of high-grade gliomas in the brain. Mol Biol Cell 29, 2509–2515. 10.1091/mbc.E18-02-0123.

3. Beeghly, G.F., Amofa, K.Y., Fischbach, C., and Kumar, S. (2022). Regulation of Tumor Invasion by the Physical Microenvironment: Lessons from Breast and Brain Cancer. Annu Rev Biomed Eng 24, 29–59. 10.1146/annurev-bioeng-110220-115419.

4. Kim, Y., Varn, F.S., Park, S.H., Yoon, B.W., Park, H.R., Lee, C., Verhaak, R.G.W., and Paek, S.H. (2021). Perspective of mesenchymal transformation in glioblastoma. Acta Neuropathol Commun 9, 50. 10.1186/s40478-021-01151-4.

5. Venturini, V., Pezzano, F., Català Castro, F., Häkkinen, H.M., Jiménez-Delgado, S., Colomer-Rosell, M., Marro, M., Tolosa-Ramon, Q., Paz-López, S., Valverde, M.A., et al. (2020). The nucleus measures shape changes for cellular proprioception to control dynamic cell behavior. Science 370. 10.1126/science.aba2644.

6. Huang, Y., Tejero, R., Lee, V.K., Brusco, C., Hannah, T., Bertucci, T.B., Junqueira Alves, C., Katsyv, I., Kluge, M., Foty, R., et al. (2021). Plexin-B2 facilitates glioblastoma infiltration by modulating cell biomechanics. Commun Biol 4, 145. 10.1038/s42003-021-01667-4.

7. Banerjee, T., Biswas, D., Pal, D.S., Miao, Y., Iglesias, P.A., and Devreotes, P.N. (2022). Spatiotemporal dynamics of membrane surface charge regulates cell polarity and migration. Nat Cell Biol 24, 1499–1515. 10.1038/s41556-022-00997-7.

8. Shinoura, N., Shamraj, O.I., Hugenholz, H., Zhu, J.G., McBlack, P., Warnick, R., Tew, J.J., Wani, M.A., and Menon, A.G. (1995). Identification and partial sequence of a cDNA that is differentially expressed in human brain tumors. Cancer Lett 89, 215–221. 030438359503690X [pii].

9. Le, A.P., Huang, Y., Pingle, S.C., Kesari, S., Wang, H., Yong, R.L., Zou, H., and Friedel, R.H. (2015). Plexin-B2 promotes invasive growth of malignant glioma. Oncotarget 6, 7293–7304. 10.18632/oncotarget.3421.

10. Buck, M. (2021). Membrane Proteins | The Plexin Family of Transmembrane Receptors. In Encyclopedia of Biological Chemistry III (Third Edition), J. Joseph, ed. (Elsevier), pp. 594–610. 10.1016/B978-0-12-819460-7.00345-5.

11. Junqueira Alves, C., Yotoko, K., Zou, H., and Friedel, R.H. (2019). Origin and evolution of plexins, semaphorins, and Met receptor tyrosine kinases. Sci Rep 9, 1970. 10.1038/s41598-019-38512-y.

12. Junqueira Alves, C., Silva Ladeira, J., Hannah, T., Pedroso Dias, R.J., Zabala Capriles, P.V., Yotoko, K., Zou, H., and Friedel, R.H. (2021). Evolution and Diversity of Semaphorins and Plexins in Choanoflagellates. Genome Biol Evol 13. 10.1093/gbe/evab035.

13. Kong, Y., Janssen, B.J., Malinauskas, T., Vangoor, V.R., Coles, C.H., Kaufmann, R., Ni, T., Gilbert, R.J., Padilla-Parra, S., Pasterkamp, R.J., and Jones, E.Y. (2016). Structural Basis for Plexin Activation and Regulation. Neuron 91, 548–560. 10.1016/j.neuron.2016.06.018.

14. Wang, Y., Pascoe, H.G., Brautigam, C.A., He, H., and Zhang, X. (2013). Structural basis for activation and non-canonical catalysis of the Rap GTPase activating protein domain of plexin. Elife 2, e01279. 10.7554/eLife.01279.

15. Fournier, A.E., Nakamura, F., Kawamoto, S., Goshima, Y., Kalb, R.G., and Strittmatter, S.M. (2000). Semaphorin3A enhances endocytosis at sites of receptor-F-actin colocalization during growth cone collapse. J Cell Biol 149, 411–422. 10.1083/jcb.149.2.411.

16. Kabayama, H., Nakamura, T., Takeuchi, M., Iwasaki, H., Taniguchi, M., Tokushige, N., and Mikoshiba, K. (2009). Ca2+ induces macropinocytosis via F-actin depolymerization during growth cone collapse. Mol Cell Neurosci 40, 27–38. 10.1016/j.mcn.2008.08.009.

17. Mehta, V., Pang, K.L., Rozbesky, D., Nather, K., Keen, A., Lachowski, D., Kong, Y., Karia, D., Ameismeier, M., Huang, J., et al. (2020). The guidance receptor plexin D1 is a mechanosensor in endothelial cells. Nature 578, 290–295. 10.1038/s41586-020-1979-4.

18. Jiang, C., Javed, A., Kaiser, L., Nava, M.M., Xu, R., Brandt, D.T., Zhao, D., Mayer, B., Fernández-Baldovinos, J., Zhou, L., et al. (2021). Mechanochemical control of epidermal stem cell divisions by B-plexins. Nat Commun 12, 1308. 10.1038/s41467-021-21513-9.

19. Junqueira Alves, C., Dariolli, R., Haydak, J., Kang, S., Hannah, T., Wiener, R.J., DeFronzo, S., Tejero, R., Gusella, G.L., Ramakrishnan, A., et al. (2021). Plexin-B2 orchestrates collective stem cell dynamics via actomyosin contractility, cytoskeletal tension and adhesion. Nat Commun 12, 6019. 10.1038/s41467-021-26296-7.

20. Tejero, R., Huang, Y., Katsyv, I., Kluge, M., Lin, J.Y., Tome-Garcia, J., Daviaud, N., Wang, Y., Zhang, B., Tsankova, N.M., et al. (2019). Gene signatures of quiescent glioblastoma cells reveal mesenchymal shift and interactions with niche microenvironment. EBioMedicine 42, 252–269. 10.1016/j.ebiom.2019.03.064.

21. Wang, G., and Galli, T. (2018). Reciprocal link between cell biomechanics and exocytosis. Traffic 19, 741–749. 10.1111/tra.12584.

22. Djakbarova, U., Madraki, Y., Chan, E.T., and Kural, C. (2021). Dynamic interplay between cell membrane tension and clathrin-mediated endocytosis. Biol Cell 113, 344–373. 10.1111/boc.202000110.

23. Gauthier, N.C., Masters, T.A., and Sheetz, M.P. (2012). Mechanical feedback between membrane tension and dynamics. Trends Cell Biol 22, 527–535. 10.1016/j.tcb.2012.07.005.

24. Dutta, D., Williamson, C.D., Cole, N.B., and Donaldson, J.G. (2012). Pitstop 2 is a potent inhibitor of clathrin-independent endocytosis. PLoS One 7, e45799. 10.1371/journal.pone.0045799.

25. Macia, E., Ehrlich, M., Massol, R., Boucrot, E., Brunner, C., and Kirchhausen, T. (2006). Dynasore, a cell-permeable inhibitor of dynamin. Dev Cell 10, 839–850. 10.1016/j.devcel.2006.04.002.

26. Masereel, B., Pochet, L., and Laeckmann, D. (2003). An overview of inhibitors of Na(+)/H(+) exchanger. Eur J Med Chem 38, 547–554. 10.1016/s0223-5234(03)00100-4.

27. Koivusalo, M., Welch, C., Hayashi, H., Scott, C.C., Kim, M., Alexander, T., Touret, N., Hahn, K.M., and Grinstein, S. (2010). Amiloride inhibits macropinocytosis by lowering submembranous pH and preventing Rac1 and Cdc42 signaling. J Cell Biol 188, 547–563. 10.1083/jcb.200908086.

28. Danylchuk, D.I., Moon, S., Xu, K., and Klymchenko, A.S. (2019). Switchable Solvatochromic Probes for Live-Cell Super-resolution Imaging of Plasma Membrane Organization. Angew Chem Int Ed Engl 58, 14920–14924. 10.1002/anie.201907690.

29. Colom, A., Derivery, E., Soleimanpour, S., Tomba, C., Molin, M.D., Sakai, N., González-Gaitán, M., Matile, S., and Roux, A. (2018). A fluorescent membrane tension probe. Nat Chem 10, 1118–1125. 10.1038/s41557-018-0127-3.

30. Fehon, R.G., McClatchey, A.I., and Bretscher, A. (2010). Organizing the cell cortex: the role of ERM proteins. Nat Rev Mol Cell Biol 11, 276–287. 10.1038/nrm2866.

31. Song, Y., Soto, J., Chen, B., Hoffman, T., Zhao, W., Zhu, N., Peng, Q., Liu, L., Ly, C., Wong, P.K., et al. (2022). Transient nuclear deformation primes epigenetic state and promotes cell reprogramming. Nat Mater 21, 1191–1199. 10.1038/s41563-022-01312-3.

32. Bera, K., Kiepas, A., Godet, I., Li, Y., Mehta, P., Ifemembi, B., Paul, C.D., Sen, A., Serra, S.A., Stoletov, K., et al. (2022). Extracellular fluid viscosity enhances cell migration and cancer dissemination. Nature 611, 365–373. 10.1038/s41586-022-05394-6.

33. Harayama, T., and Riezman, H. (2018). Understanding the diversity of membrane lipid composition. Nat Rev Mol Cell Biol 19, 281–296. 10.1038/nrm.2017.138.

34. Balla, T., and Várnai, P. (2009). Visualization of cellular phosphoinositide pools with GFP-fused protein-domains. Curr Protoc Cell Biol Chapter 24, Unit 24.24. 10.1002/0471143030.cb2404s42.

35. Várnai, P., and Balla, T. (1998). Visualization of phosphoinositides that bind pleckstrin homology domains: calcium- and agonist-induced dynamic changes and relationship to myo-[3H]inositol-labeled phosphoinositide pools. J Cell Biol 143, 501–510. 10.1083/jcb.143.2.501.

36. Dickson, E.J., and Hille, B. (2019). Understanding phosphoinositides: rare, dynamic, and essential membrane phospholipids. Biochem J 476, 1–23. 10.1042/BCJ20180022.

37. Coxon, K.M., Duggan, J., Cordeiro, M.F., and Moss, S.E. (2011). Purification of annexin V and its use in the detection of apoptotic cells. Methods Mol Biol 731, 293–308. 10.1007/978-1-61779-080-5_24.

38. McLaughlin, S., Wang, J., Gambhir, A., and Murray, D. (2002). PIP(2) and proteins: interactions, organization, and information flow. Annu Rev Biophys Biomol Struct 31, 151–175. 10.1146/annurev.biophys.31.082901.134259.

39. Ma, Y., Poole, K., Goyette, J., and Gaus, K. (2017). Introducing Membrane Charge and Membrane Potential to T Cell Signaling. Front Immunol 8, 1513. 10.3389/fimmu.2017.01513.

40. Yeung, T., Terebiznik, M., Yu, L., Silvius, J., Abidi, W.M., Philips, M., Levine, T., Kapus, A., and Grinstein, S. (2006). Receptor activation alters inner surface potential during phagocytosis. Science 313, 347–351. 10.1126/science.1129551.

41. Miller, E.W., Lin, J.Y., Frady, E.P., Steinbach, P.A., Kristan, W.B., and Tsien, R.Y. (2012). Optically monitoring voltage in neurons by photo-induced electron transfer through molecular wires. Proc Natl Acad Sci U S A 109, 2114–2119. 10.1073/pnas.1120694109.

42. Pascoe, H.G., Wang, Y., and Zhang, X. (2015). Structural mechanisms of plexin signaling. Prog Biophys Mol Biol 118, 161–168. 10.1016/j.pbiomolbio.2015.03.006.

43. Kooistra, M.R., Dubé, N., and Bos, J.L. (2007). Rap1: a key regulator in cell-cell junction formation. J Cell Sci 120, 17–22. 10.1242/jcs.03306.

44. Bos, J.L. (2018). From Ras to Rap and Back, a Journey of 35 Years. Cold Spring Harb Perspect Med 8. 10.1101/cshperspect.a031468.

45. Lomakin, A.J., Cattin, C.J., Cuvelier, D., Alraies, Z., Molina, M., Nader, G.P.F., Srivastava, N., Sáez, P.J., Garcia-Arcos, J.M., Zhitnyak, I.Y., et al. (2020). The nucleus acts as a ruler tailoring cell responses to spatial constraints. Science 370. 10.1126/science.aba2894.

46. Lanzetti, L., and Di Fiore, P.P. (2008). Endocytosis and cancer: an ‘insider’ network with dangerous liaisons. Traffic 9, 2011–2021. 10.1111/j.1600-0854.2008.00816.x.

47. Khan, I., and Steeg, P.S. (2021). Endocytosis: a pivotal pathway for regulating metastasis. Br J Cancer 124, 66–75. 10.1038/s41416-020-01179-8.

48. Tojima, T., Itofusa, R., and Kamiguchi, H. (2010). Asymmetric clathrin-mediated endocytosis drives repulsive growth cone guidance. Neuron 66, 370–377. 10.1016/j.neuron.2010.04.007.

49. Jurney, W.M., Gallo, G., Letourneau, P.C., and McLoon, S.C. (2002). Rac1-mediated endocytosis during ephrin-A2- and semaphorin 3A-induced growth cone collapse. J Neurosci 22, 6019–6028. 10.1523/JNEUROSCI.22-14-06019.2002.

50. Wu, K.Y., He, M., Hou, Q.Q., Sheng, A.L., Yuan, L., Liu, F., Liu, W.W., Li, G., Jiang, X.Y., and Luo, Z.G. (2014). Semaphorin 3A activates the guanosine triphosphatase Rab5 to promote growth cone collapse and organize callosal axon projections. Sci Signal 7, ra81. 10.1126/scisignal.2005334.

51. Tanaka, H., Kanatome, A., and Takagi, S. (2020). Involvement of the synaptotagmin/stonin2 system in vesicular transport regulated by semaphorins in Caenorhabditis elegans epidermal cells. Genes Cells 25, 391–401. 10.1111/gtc.12765.

52. Goldenberg, N.M., and Steinberg, B.E. (2010). Surface charge: a key determinant of protein localization and function. Cancer Res 70, 1277–1280. 10.1158/0008-5472.CAN-09-2905.

53. Heo, W.D., Inoue, T., Park, W.S., Kim, M.L., Park, B.O., Wandless, T.J., and Meyer, T. (2006). PI(3,4,5)P3 and PI(4,5)P2 lipids target proteins with polybasic clusters to the plasma membrane. Science 314, 1458–1461. 10.1126/science.1134389.

54. Suh, B.C., and Hille, B. (2007). Electrostatic interaction of internal Mg2+ with membrane PIP2 Seen with KCNQ K+ channels. J Gen Physiol 130, 241–256. 10.1085/jgp.200709821.

55. Worzfeld, T., Swiercz, J.M., Senturk, A., Genz, B., Korostylev, A., Deng, S., Xia, J., Hoshino, M., Epstein, J.A., Chan, A.M., et al. (2014). Genetic dissection of plexin signaling in vivo. Proc Natl Acad Sci U S A 111, 2194–2199. 10.1073/pnas.1308418111.

56. Hota, P.K., and Buck, M. (2012). Plexin structures are coming: opportunities for multilevel investigations of semaphorin guidance receptors, their cell signaling mechanisms, and functions. Cell Mol Life Sci 69, 3765–3805. 10.1007/s00018-012-1019-0.

57. Wang, Q., Hu, B., Hu, X., Kim, H., Squatrito, M., Scarpace, L., deCarvalho, A.C., Lyu, S., Li, P., Li, Y., et al. (2017). Tumor Evolution of Glioma-Intrinsic Gene Expression Subtypes Associates with Immunological Changes in the Microenvironment. Cancer Cell 32, 42–56.e46. 10.1016/j.ccell.2017.06.003.

58. Burk, K., Mire, E., Bellon, A., Hocine, M., Guillot, J., Moraes, F., Yoshida, Y., Simons, M., Chauvet, S., and Mann, F. (2017). Post-endocytic sorting of Plexin-D1 controls signal transduction and development of axonal and vascular circuits. Nat Commun 8, 14508. 10.1038/ncomms14508.

59. Jayashankar, V., and Edinger, A.L. (2020). Macropinocytosis confers resistance to therapies targeting cancer anabolism. Nat Commun 11, 1121. 10.1038/s41467-020-14928-3.

60. Bodhinathan, K., and Slesinger, P.A. (2013). Molecular mechanism underlying ethanol activation of G-protein-gated inwardly rectifying potassium channels. Proc Natl Acad Sci U S A 110, 18309–18314. 10.1073/pnas.1311406110.

61. Rosenson, R.S., McCormick, A., and Uretz, E.F. (1996). Distribution of blood viscosity values and biochemical correlates in healthy adults. Clin Chem 42, 1189–1195.

