## Supplementary material for "Invasion of glioma cells through confined space requires membrane tension regulation and mechano-electrical coupling via Plexin-B2": Suppl Figs and Movie legends

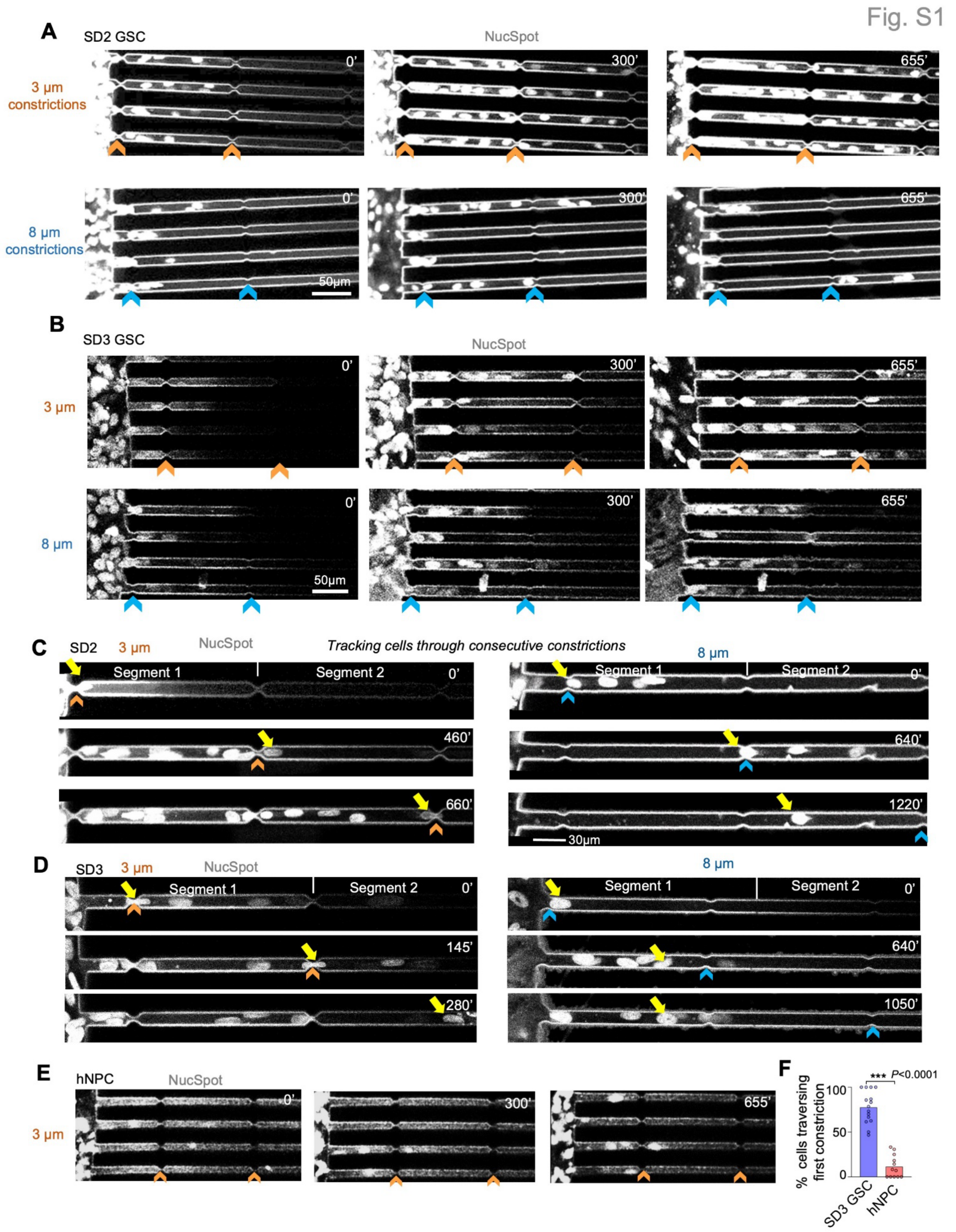


**Figure S1. Patient-derived GSCs are adept at confined migration, related to Figure 1.**

(A, B) Additional still frames from live-cell videography show that SD2 (A) and SD3 (B) GSCs labeled with

NucSpot pass through microchannels with 3 µm constrictions (orange chevrons) more efficiently than 8 µm constrictions (blue chevrons). Time stamps in minutes are in relation to start of imaging.

(C, D) Tracking of individual SD2 or SD3 GSC (yellow arrows) traversing consecutive constrictions (chevrons). Passage through the first 3 µm constriction led to accelerated passage through the second one. Note longer time frames for cells traversing the 8 µm constrictions.

(E) Still frames from live-cell videography show slower migration of hNPCs in the microchannels with 3 µm constrictions as compared to GSCs, with only few hNPCs reaching the second constriction by 655 minutes (compared to **Fig. S1A and B**, 3 µm constrictions).

(F) Bar graphs showing the percentage of SD3 GSCs and hNPCs that had successfully traversed the first 3 µm constriction upon entering the first segment of microchannels within the timeframe. Each data point represents cells in one microchannel. For SD3 GSC, n=170 cells from 15 channels; for hNPCs, n=132 cells from 12 channels. Data show mean ± SEM. Two-sided unpaired t-test. Related to **movie S3**.

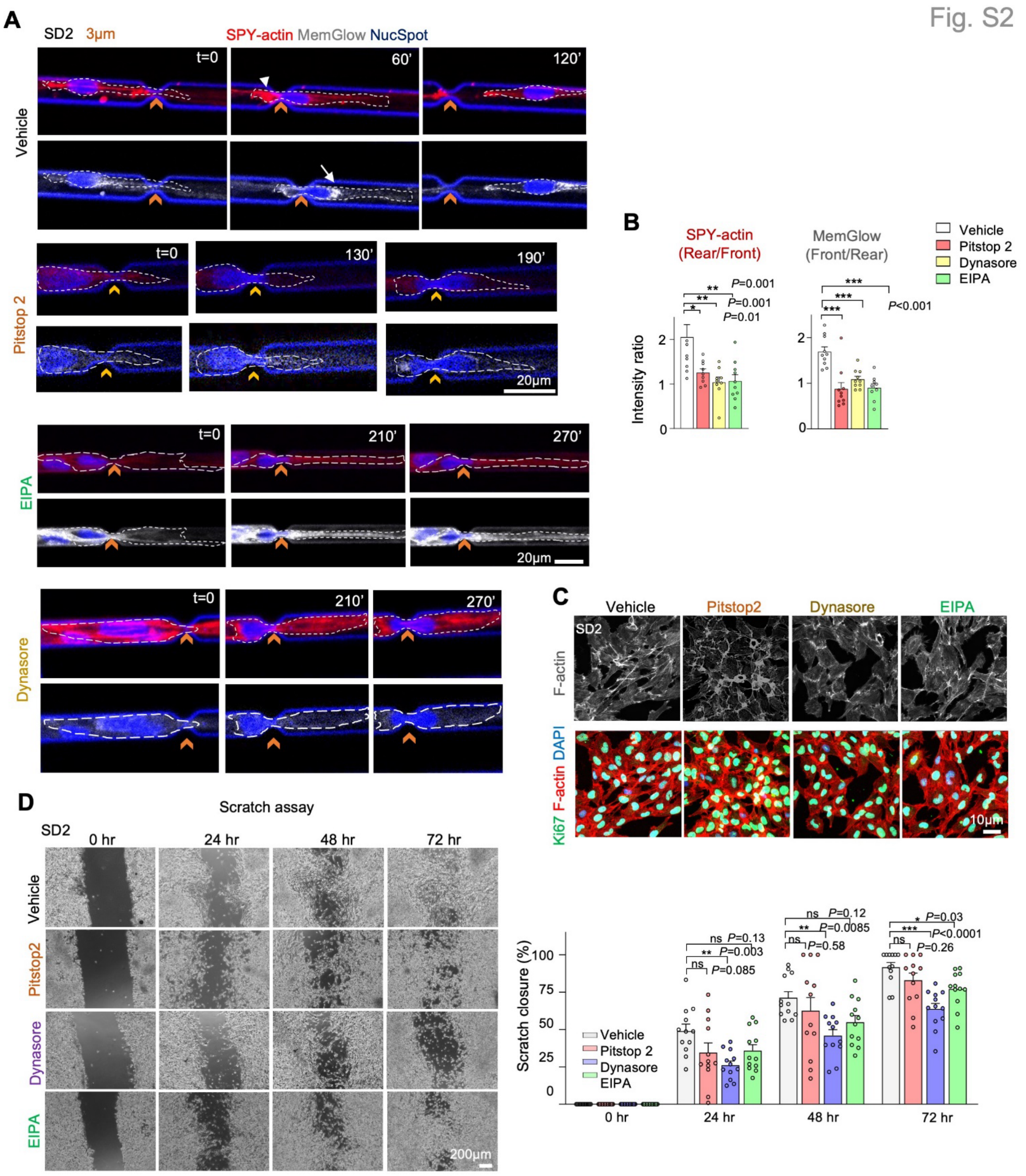


**Figure S2. Endocytosis inhibitors block confined migration of GSCs, related to Figure 1.**

(A) Still frames from live-cell videography show that migration of SD2 GSCs in microchannels with 3 μm constrictions (orange chevrons) was impaired by endocytosis inhibitors, with disruption of accumulation of SPY-actin at rear (arrowhead) and MemGlow at cell front (arrow) seen in vehicle-treated cells.

(B) Quantifications of fluorescence intensity ratio of SPY-actin or MemGlow at rear vs. front during confined migration of SD2 GSCs treated with vehicle or endocytosis inhibitors. n= 9-10 cells for each condition. One-way ANOVA followed by Dunnett's multiple comparisons test. Data represent mean ± SEM.

(C) IF images of SD2 GSCs in 2D cultures show no major effect on F-actin (phalloidin) or proliferation (Ki67) by endocytosis inhibitors. DAPI for nuclear staining.

(D) Images and quantifications of scratch wound assay show overall similar wound closure performance of SD2 GSCs treated with vehicle or endocytosis inhibitors, except for a modest delay by Dynasore (24-72 hr) or EIPA (only at 72 hr). Bar graphs show the extent of scratch closure in comparison to 0 hr. n=5 fields/group from 3 independent replicates. One-way ANOVA followed by Dunnett's multiple comparisons test. Data represent mean ± SEM.

Fig. S3

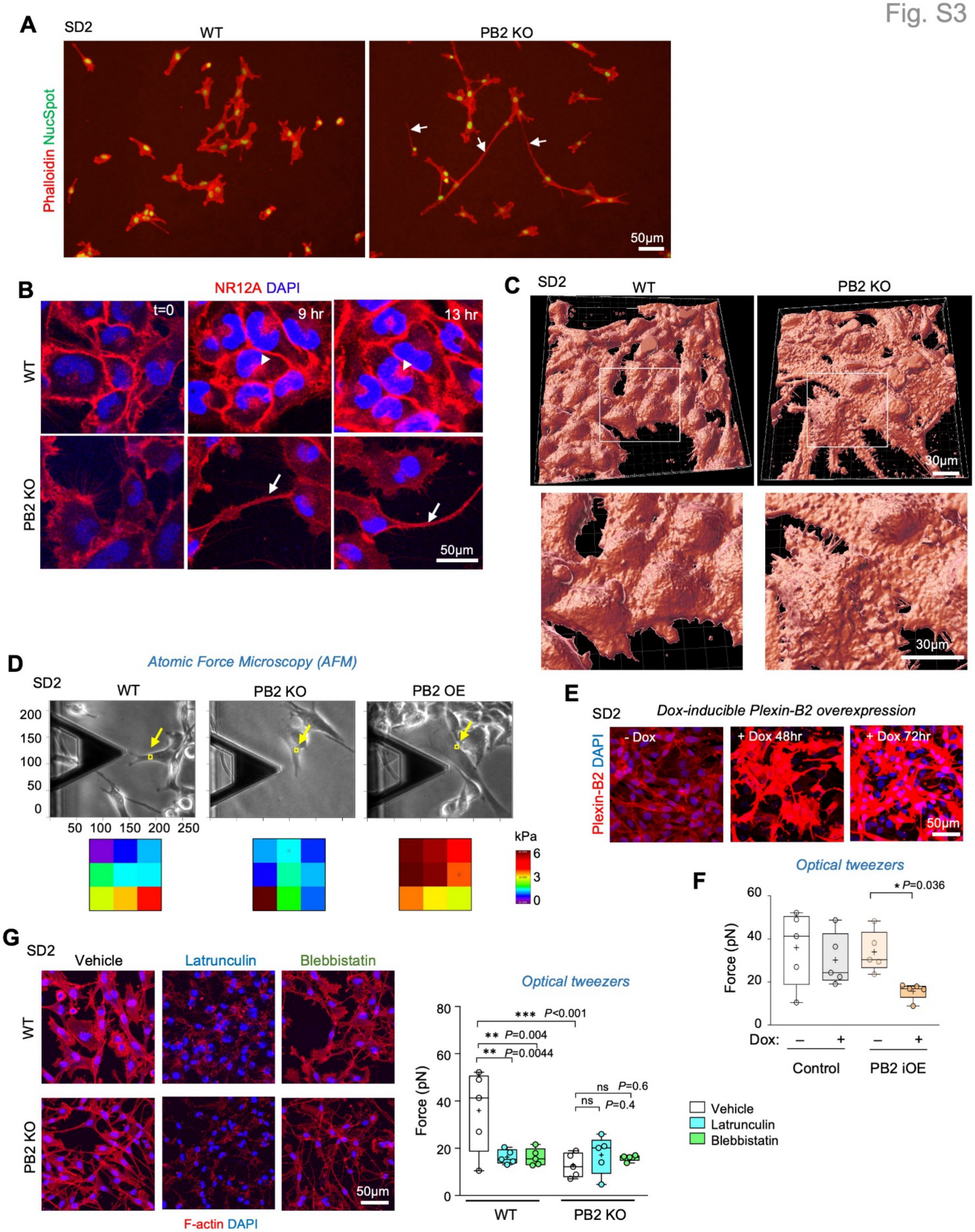


**Figure S3. Plexin-B2 regulates cytoskeletal stiffness and membrane tension in GSCs, related to Figure 2.**

(A) IF images of SD2 GSCs show long processes from PB2 KO cells (arrows) containing F-actin (Alexa 568 phalloidin), which were not detected in WT cells. NucSpot for nuclear staining.

(B) Still frames from live-cell imaging of SD2 GSCs labeled by Nile red (NR12A, a solvatochromic plasma membrane-targeting dye) show reduced NR12$^+$ endosome in PB2 KO compared to WT GSCs (arrowheads). Arrows point to long processes of PB2 KO cells. NucSpot for nuclear staining.

(C) Top panels, 3D surface rendering of GSCs labeled with Nile red (NR12A) dye using IMARIS software (Bitplane). Bottom panels are enlarged images of boxed areas above, showing more pronounced cell membrane ruffling or wrinkles in PB2 KO relative to WT cells.

(D) Top panels, phase-contrast images show the area of GSC (yellow box, arrow) measured by atomic force microscopy (AFM) elastography. Bottom panels, corresponding AFM stiffness heatmaps of boxed area above.

(E) Top, IF images show induced overexpression of Plexin-B2 in SD2 GSCs after 48 and 72 hr of Dox induction. DAPI for nuclear staining.

(F) Quantifications of membrane tension measured by optical tweezers of SD2 GSCs with or without Dox induction (72 hr) for control or PB2 iOE. Box plots show 25–75% quartiles, median (line), mean (plus sign). Data collected from 5 cells per group. Kruskal–Wallis test followed by Dunn's multiple comparisons test.

(G) Left, IF images of WT or PB2 KO SD2 GSCs, stained for phalloidin and DAPI after treatment with vehicle or actomyosin inhibitors latrunculin or blebbistatin. Right, box plots of membrane tension measured with optical tweezers, 25–75% quartiles, median (line), mean (plus sign). Data collected from 5 cells per group. One-way ANOVA followed by Dunnett's multiple comparisons test.

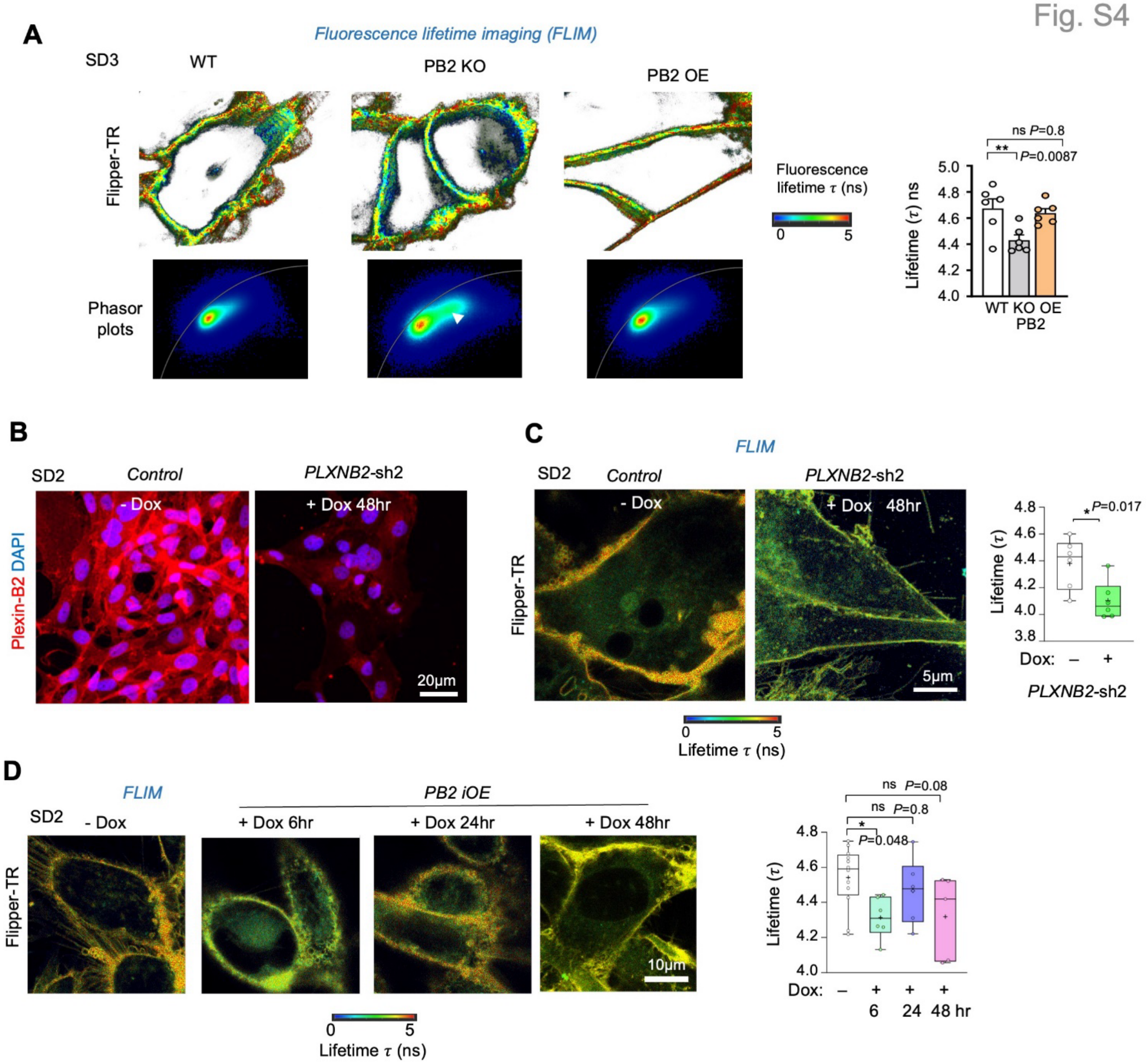


**Figure S4. Plexin-B2 regulates membrane tension in GSCs measured by Flipper-TR dye, related to Figure 2.**

(A) Top, representative fluorescence lifetime images (FLIM) of SD3 GSCs labeled with the Flipper-TR membrane dye. Heatmap of lifetime values is shown on right. Bottom, phasor plots of lifetime data from FLIM, with arrowhead indicating a shift to shorter lifetime values for PB2 KO cell. Right, bar graphs of average fluorescence lifetime. n=6 images per genotype. One-way ANOVA followed by Dunnett's multiple comparisons test. Data represent mean ± SEM.

(B) IF images of SD2 GSCs show Dox-induced PB2 knockdown (*PLXNB2*-sh2) after 48 hr. DAPI for nuclear staining.

(C) Left, FLIM of SD2 GSCs labeled byFlipper-TR. Right, box plots show average fluorescence lifetime, with 25–75% quartiles, median (line), mean (plus sign). n= 6 cells per group. Two-sided unpaired t-test.

(D) Left, FLIM of SD2 GSCs at different time points with or without Dox-inducible PB2 OE. Right, box plots of average fluorescence lifetime, with 25–75% quartiles, median (line), and mean (plus sign). n=12 cells for -Dox, n=5-6 cells for +Dox groups. One-way ANOVA followed by Dunnett's multiple comparisons test. Data represent mean ± SEM.

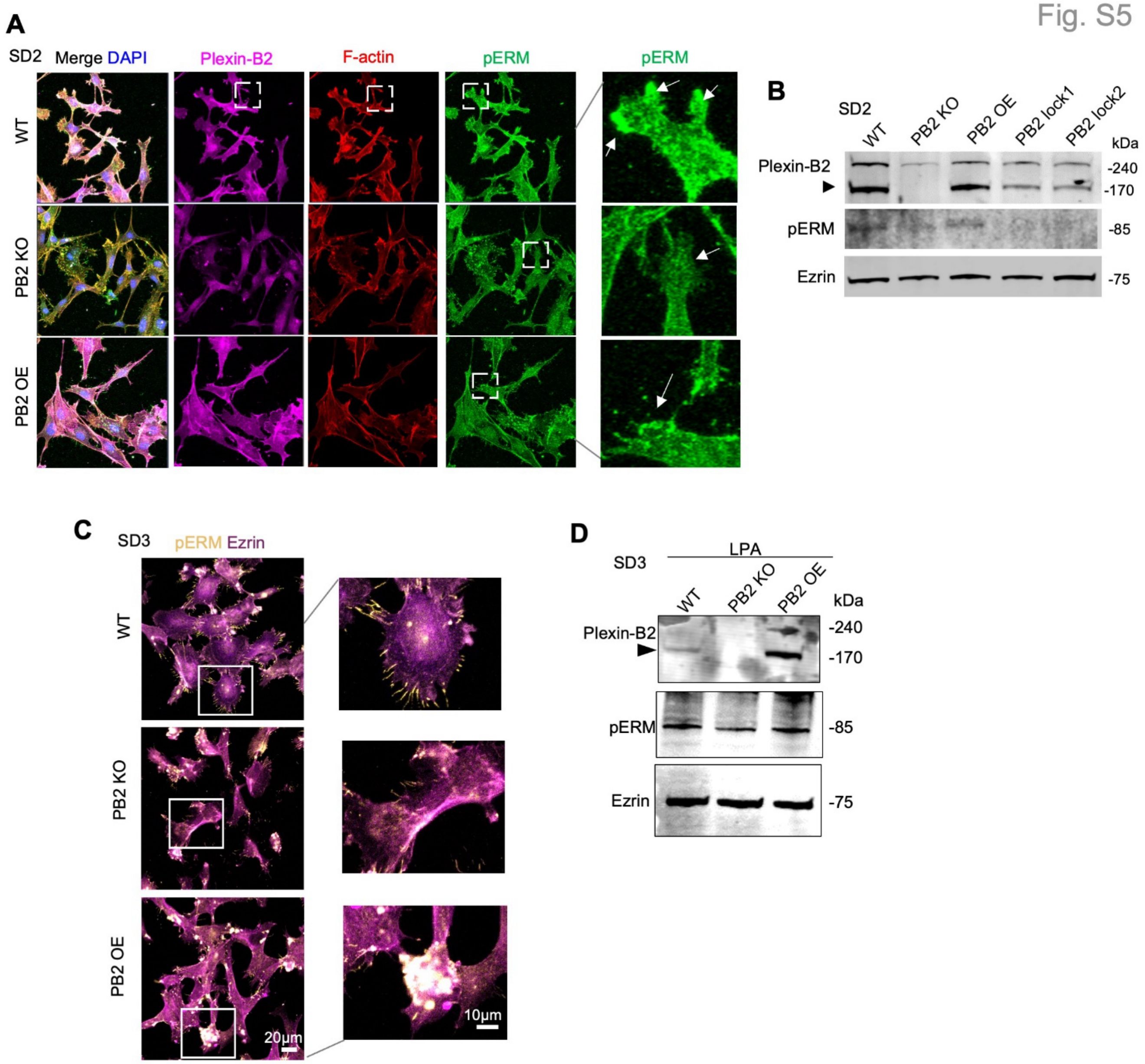


**Figure S5. Plexin-B2 affects pERM localization in GSCs, related to Figure 2.**

(A) Confocal IF images of SD2 GSCs stained for pERM (phospho-Ezrin/Radixin/Moesin), Plexin-B2, F-actin (Alexa 568 phalloidin), and nuclear staining (DAPI). Enlarged images of boxed areas on the right show high pERM signals in WT cells, particularly at the edge of lamellipodia (arrows) relative to PB2 KO or OE cells.

(B) Western blots show expression of Plexin-B2, pERM, and total Ezrin in SD2 GSCs. Note reduced pERM levels in GSCs with PB2 KO or lock mutants compared to WT GSCs. Also note re-expression of PB2 lock1 and lock2 mutants for rescue experiments (see Fig. 7).

(C) Confocal IF images of SD3 GSCs stained for pERM and total Ezrin. Enlarged images of boxed areas on the right show localized pERM signals at tip of cell processes of WT cell, which showed reduction in PB2 KO and abnormal aggregations in PB2 OE GSCs.

(D) Western blots show expression of Plexin-B2, pERM, and total Ezrin in SD3 GSCs treated with oleoyl lysophosphatidic acid (LPA; 5 μM) for 5 min to stimulate pERM expression. Note reduced pERM levels in PB2 KO and a slight increase in PB2 OE cells relative to WT GSCs.

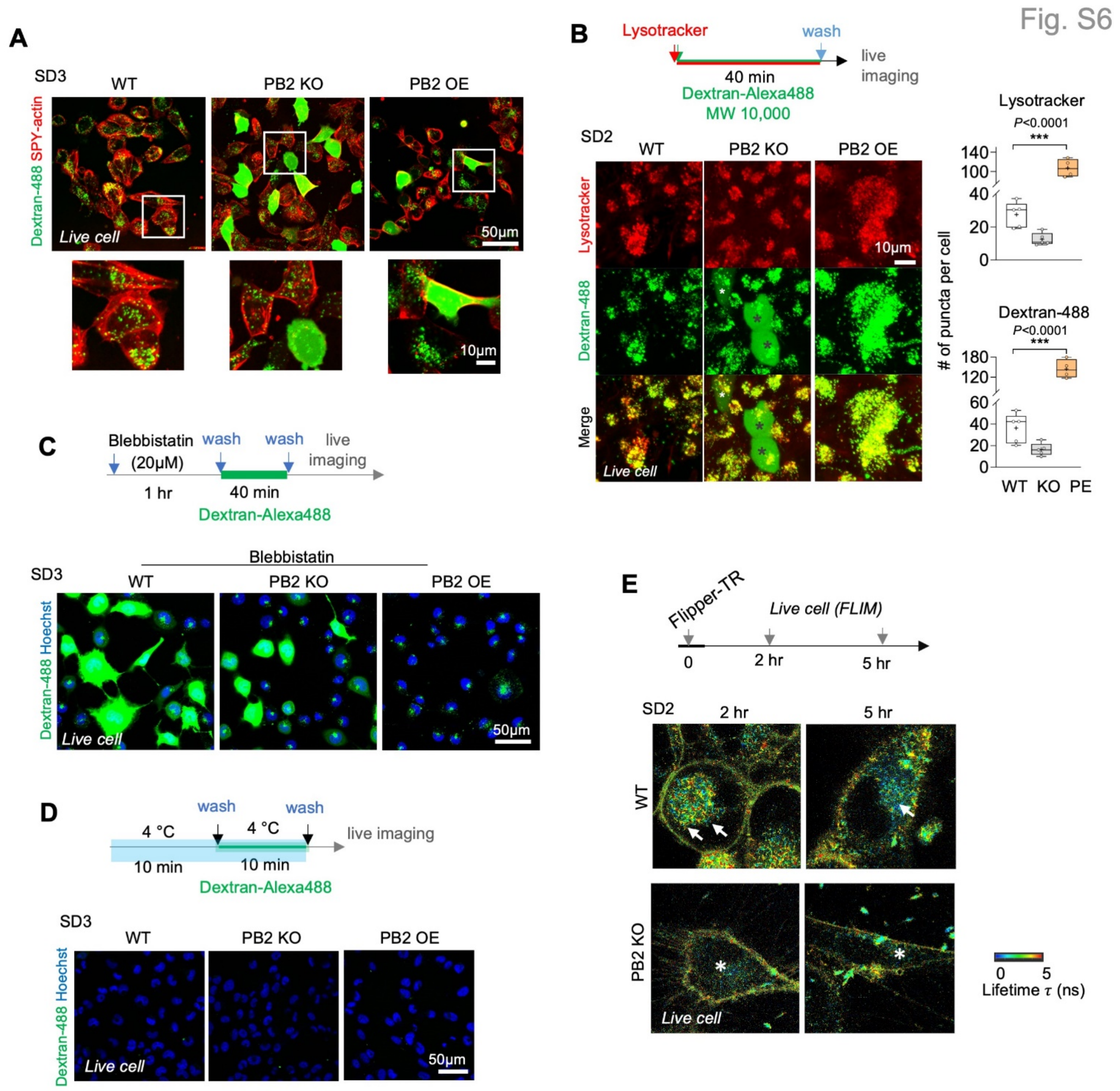


**Figure S6. Plexin-B2 affects membrane permeability and membrane internalization in GSCs, related to Figure 3.**

(A) Live-cell imaging of SD3 GSCs exposed to dextran-Alexa488 for 40 min and labeled with SPY-actin. Enlarged images of boxed areas below highlight that some of PB2 KO and OE cells harbor diffuse cytoplastic dextran fluorescence. PB2 KO GSCs also contained less dextran$^+$ puncta (endosomes) than WT.

(B) Top, experimental scheme for dextran endocytosis assay co-labeled with Lysotracker dye. Bottom, live cell images of GSCs show co-localization of Lysotracker with dextran puncta, but not with the diffuse cytoplasmic dextran (asterisks). Right, box plots show the number of dextran and Lysotracker positive dots per cell, with 25–75% quantiles, minimal and maximal values (whiskers), median (line), and mean (cross). n=4-5 cells/genotype. One-way ANOVA followed by Dunnett's multiple comparisons test.

(C) Top, dextran endocytosis assay of SD3 GSCs pre-treated with myosin inhibitor blebbistatin (20 μM). Bottom, live cell imaging shows appearance of diffuse cytoplasmic dextran-Alexa488 in WT SD3 treated with blebbistatin, phenocopying PB2 KO, while PB2 OE cells did not display cytoplasmic dextran leakage. Hoechst for nuclear visualization.

(D) Top, experimental scheme for dextran endocytosis assay of SD3 GSCs maintained at 4°C. Bottom, confocal live images show that 4°C culture blocked the cytoplasmic dextran fluorescence in GSCs of all genotypes.

(E) Top, experimental scheme for live-cell imaging of GSCs after 2 or 5 hr incubation with Flipper-TR. Bottom, FLIM show higher number of endosomes (arrows) in WT cells relative to PB2 KO cells (asterisks). Heatmap indicates lifetime values. Note that the Flipper-TR lifetime is typically lower on endosomes than on plasma membrane.

Fig. S7

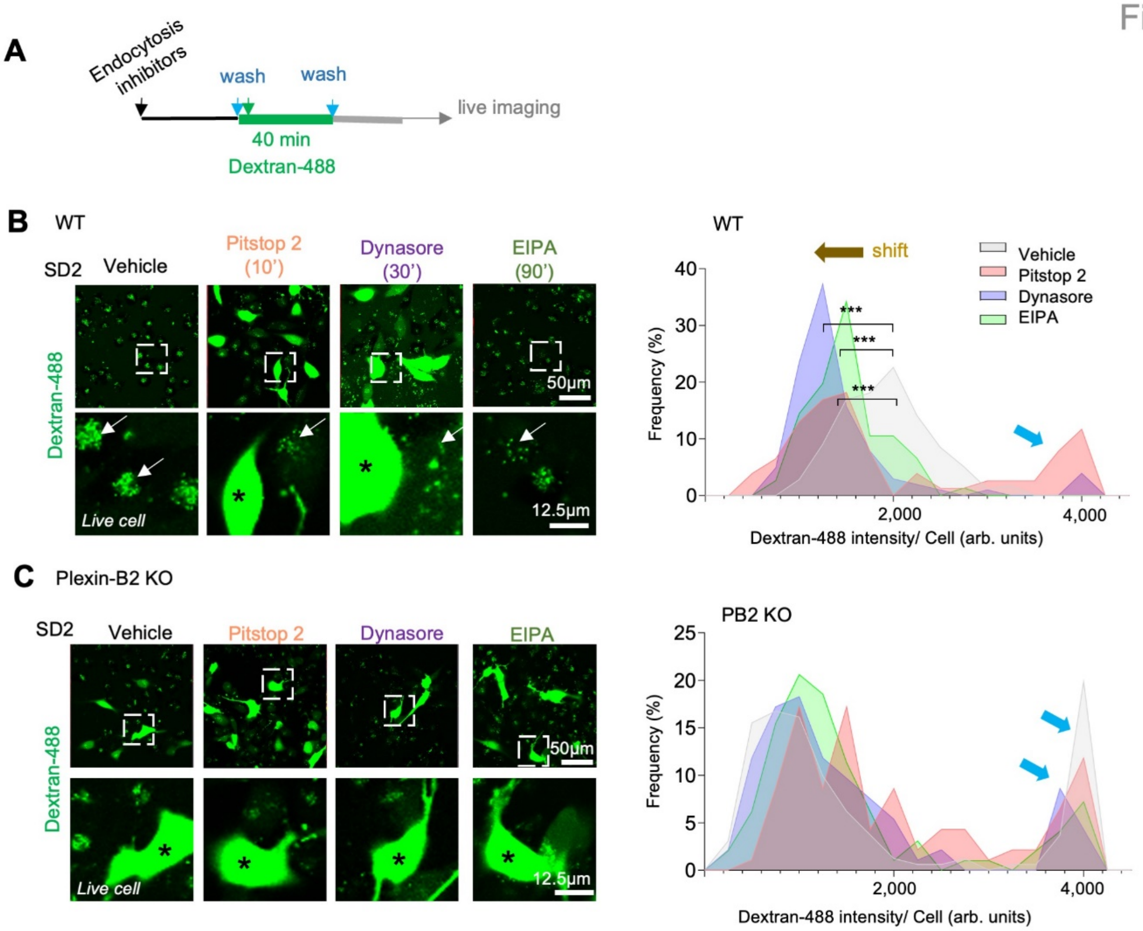


**Figure S7. Endocytosis inhibitors and Plexin-B2 KO increase membrane permeability, related to Figure 3.**

(A) Experimental scheme of dextran uptake assay.

(B, C) Left, confocal live-cell images show that endocytosis inhibitors reduced dextran puncta (arrow) but increased diffuse cytoplastic dextran (asterisk) in WT SD2 GSCs (B), but the inhibitors had no further effect on PB2 KO cells (C). Right, histograms show reduced dextra punta (denoted by leftward shift, brown arrow), and appearance of bimodal distribution of dextran-Alexa 488 fluorescence (blue arrows) by endocytosis inhibitors or PB2 KO. For WT GSCs: vehicle, n=177 cells; Pitstop 2, n= 77; Dynasore, n=102; and EIPA, n=76. For PB2 KO: vehicle, n=161 cells; Pitstop 2, n= 93; Dynasore, n=93; and EIPA, n=97. One-way ANOVA followed by Dunnett's multiple comparisons test.

Fig. S8

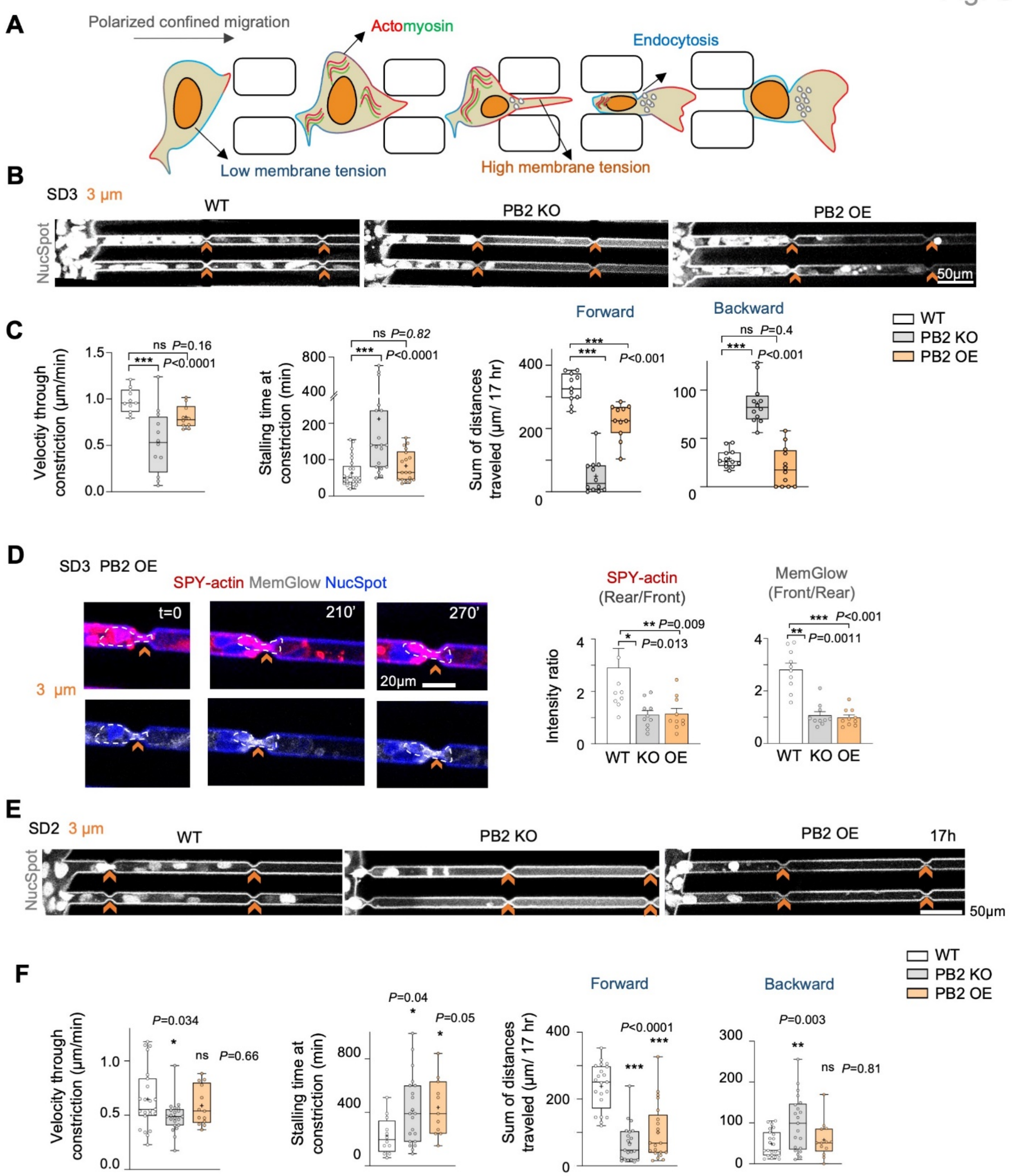


**Figure S8. Plexin-B2 manipulation compromises confined migration of GSCs, related to Figure 4.**

(A) Scheme depiction of alteration of membrane tension and regionalized endocytosis at cell front and F-actin at cell rear during polarized confined migration of GSCs.

(B) Still frames from live-cell imaging of NucSpot-labeled GSCs show reduced migration of PB2 KO or OE SD3 GSCs in microchannels with 3 µm constriction (orange chevrons) as compared to WT.

(C) Quantifications of velocity through constrictions, stalling time at constrictions, sums of distances traveled forward or backward over 17 hr. Box plots show 25–75% quantiles, minimal and maximal values (whiskers), median (line), and mean (cross). Of note, WT and PB2 KO quantifications are shown in Fig. 4B. For stalling time at constrictions: WT, n=25 cells; PB2 KO, n=19, and PB2 OE, n=18. For the rest, n=10-12 cells. One-way ANOVA followed by Dunnett's multiple comparisons test.

(D) Left, still frames from live-cell imaging of PB2 OE SD3 GSCs stalled at 3 μm constrictions. Dashed lines delineate cell boundary. Note disorganized F-actin (SPY555-actin) and MemGlow-labeled endosomes without regionalization. Right, bar graphs showing the ratio of fluorescence intensity for SPY-actin or MemGlow at rear vs. front of cells during confined migration. Data represent mean ± SEM. Of note, WT and PB2 KO quantifications are the same as in Fig. 4B. n=10 cells per group. Kruskal–Wallis test followed by Dunn's multiple comparisons test.

(E) Still frames from live-cell imaging of NucSpot-labeled SD2 GSCs shows reduced migration of PB2 KO or OE cells into the microchannel with 3 μm constrictions as compared to WT.

(F) Quantifications of velocity through constrictions, stalling time at constrictions, sums of distances traveled for forward or backward movement over 17 hr. For velocity through constriction and sum of backward movement: WT, n=23 cells; PB2 KO, n=21, and PB2 OE, n=13. For stalling time at the constriction: WT, n=12 cells; PB2 KO, n=21, and PB2 OE, n=11. For sum of forward movement: n=21 each. One-way ANOVA followed by Dunnett's multiple comparisons test.

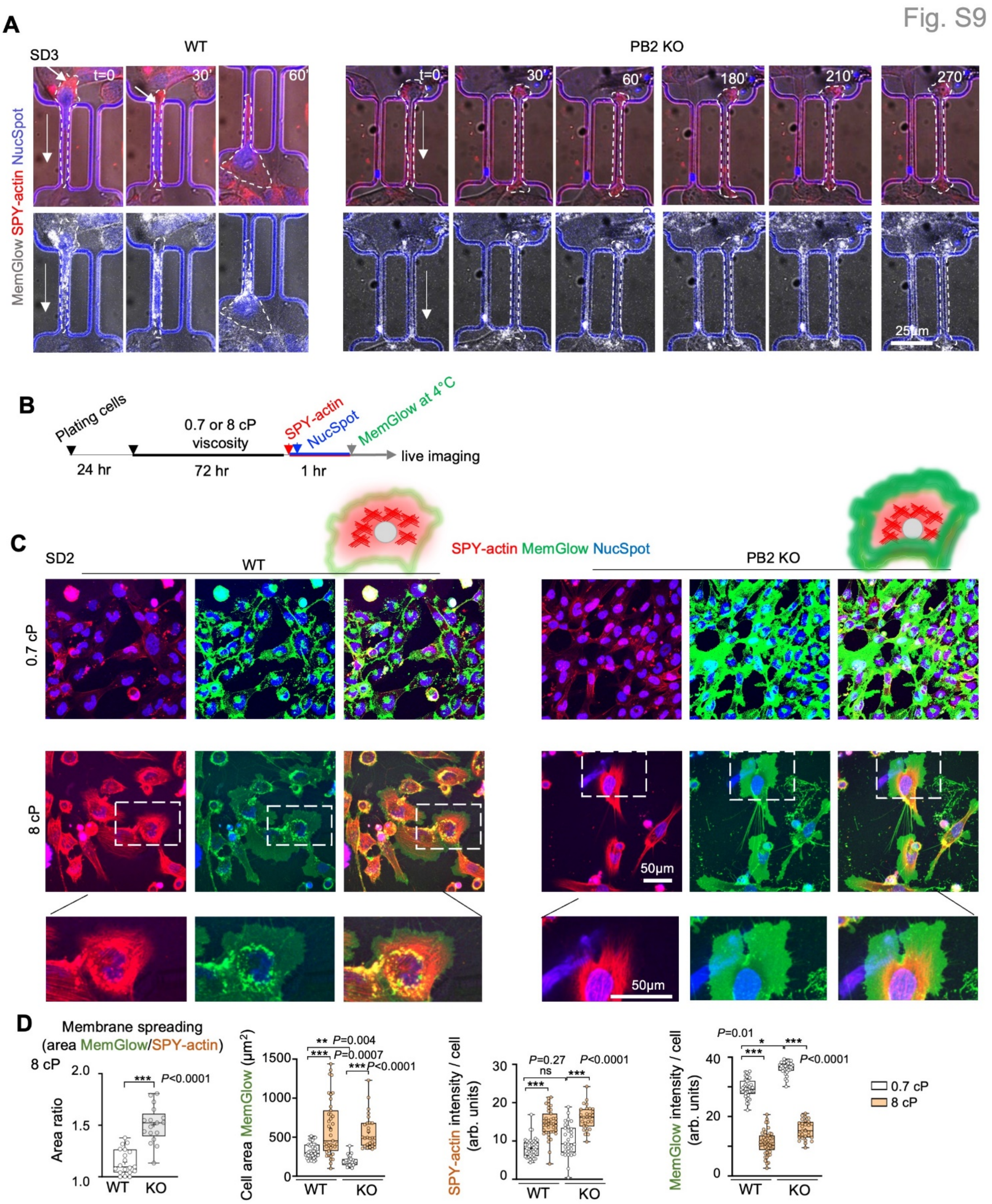


**Figure S9. Viscosity induced actin contraction is independent of Plexin-B2, related to Figure 4.**

(A) Additional still images from videography show that WT SD3 GSC traversed the 3 μm-wide tunnel within 60 min, but PB2 KO cell was stalled for at least 270 min. Long arrows point to migration direction. Note assembly of F-actin (arrow) at the rear of WT but not PB2 KO GSC, while MemGlow-labeled endosomes appeared throughout WT cells, but reduced in PB2 KO cells. Dashed lines delineate cell boundary.

(B) Experimental scheme of increasing viscosity in culture media, followed by live cell imaging.

(C) Confocal live cell imaging shows increased cortical actin (SPY555-actin) in response to higher viscosity in both WT and PB2 KO cells, but the membrane spread (MemGlow) was more pronounced in KO than WT cells (cartoon illustration on top; enlarged images of boxed area at bottom).

(D) Quantifications of membrane/cortical actin membrane spreading ratio, cell area, and SPY-actin and MemGlow fluorescent intensities in SD2 GSCs in response to high vs. low viscosity media (8 vs. 0.7 cP) for 72 hr. Box plots show 25–75% quantiles, minimal and maximal values (whiskers), median (line), and mean (cross). For membrane/actin spread ratio: n=17-19 cells/genotype, two-sided unpaired t-test. For the rest, n=28-39 cells per condition, Kruskal–Wallis test followed by Dunn's multiple comparisons test.

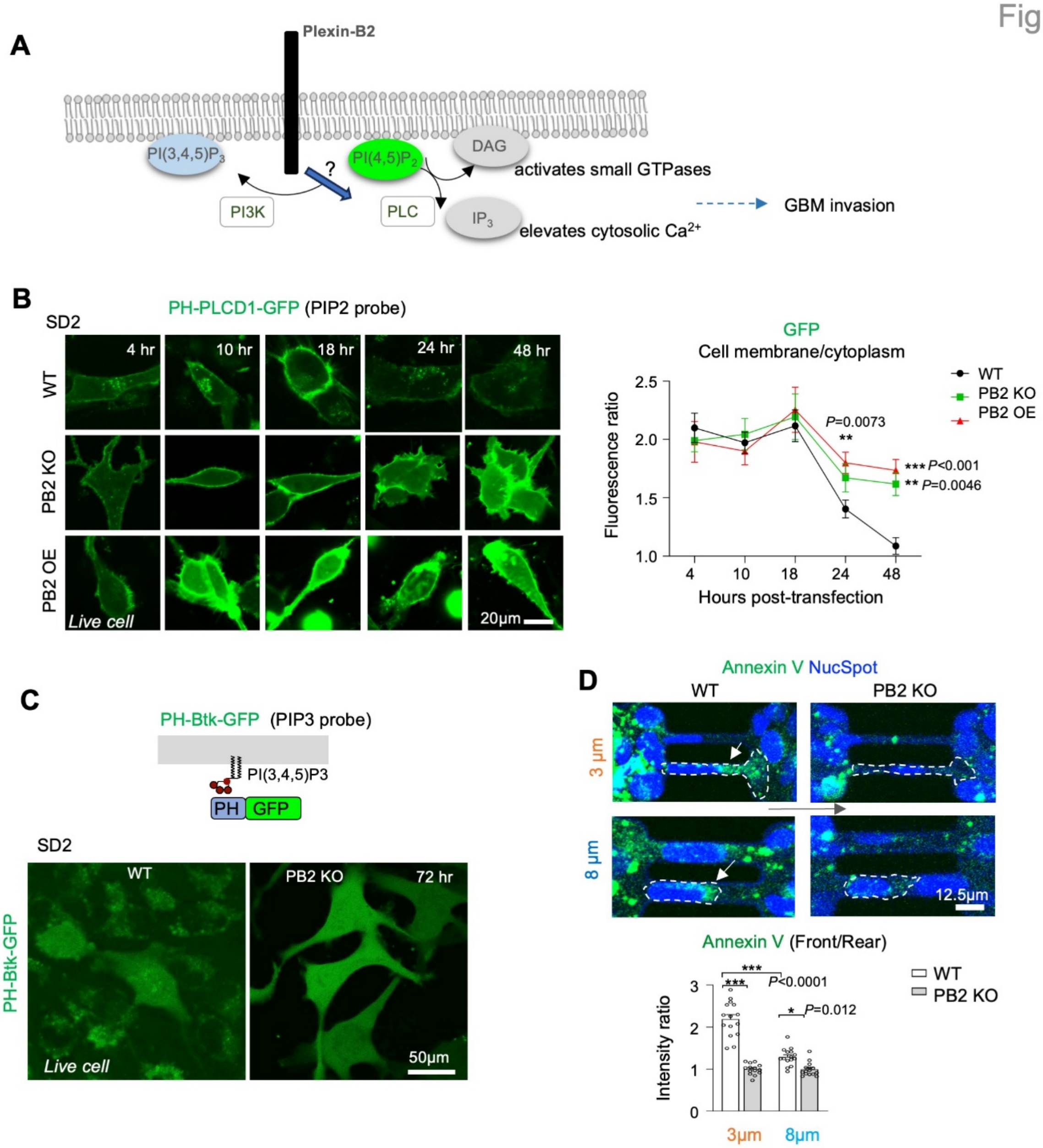


**Figure S10. Plexin-B2 affects anionic membrane lipid composition, related to Figure 6.**

(A) Diagram of regulation of PI(4,5)$P_2$ by PLC and PI3K and a potential effect from Plexin-B2-mediated membrane internalization/endocytosis during polarized migration of GBM cells.

(B) Live fluorescence imaging of SD GSCs expressing PIP2 probe (PH(PLCD1)-GFP) at 4 to 48 hr after transfection. Note the appearance of membrane GFP signals at 18 hr in WT cells followed by internalization, but sustained membrane GFP signals in PB2 KO and OE cells. Quantifications show the ratio of membrane vs. internalized GFP signals. For 4 and 10 hr: n=15 cells for each genotype. For 18 hr: n=27 for WT, n=15 for PB2 KO or OE. For 24 hr: n=30 for WT or OE, n=15 for PB2 KO. For 48 hr: n=24 for WT, n=15 for PB2 KO or OE. Two-way ANOVA followed by Dunnett's post hoc test.

(C) Top, diagram of PIP3 probe (PH(Btk)-GFP). Bottom, live cell imaging shows low GFP levels in both WT and PB2 KO GSCs.

(D) Still images from videography show accumulation of annexin V (binding to phosphatidylserine (PS)) at cell front (arrows) of WT GSCs traversing tunnels, more so in the 3 μm than the 8 μm tunnel, but absent in PB2 KO cells. Dashed lines delineate cell boundary. Long arrows denote the direction of migration. Bottom, quantifications show ratio of annexin V intensity at front vs. rear during passage through tunnels. n=14-15 cells/condition. One-way ANOVA followed by Tukey's multiple comparison test. Data represent mean ± SEM.

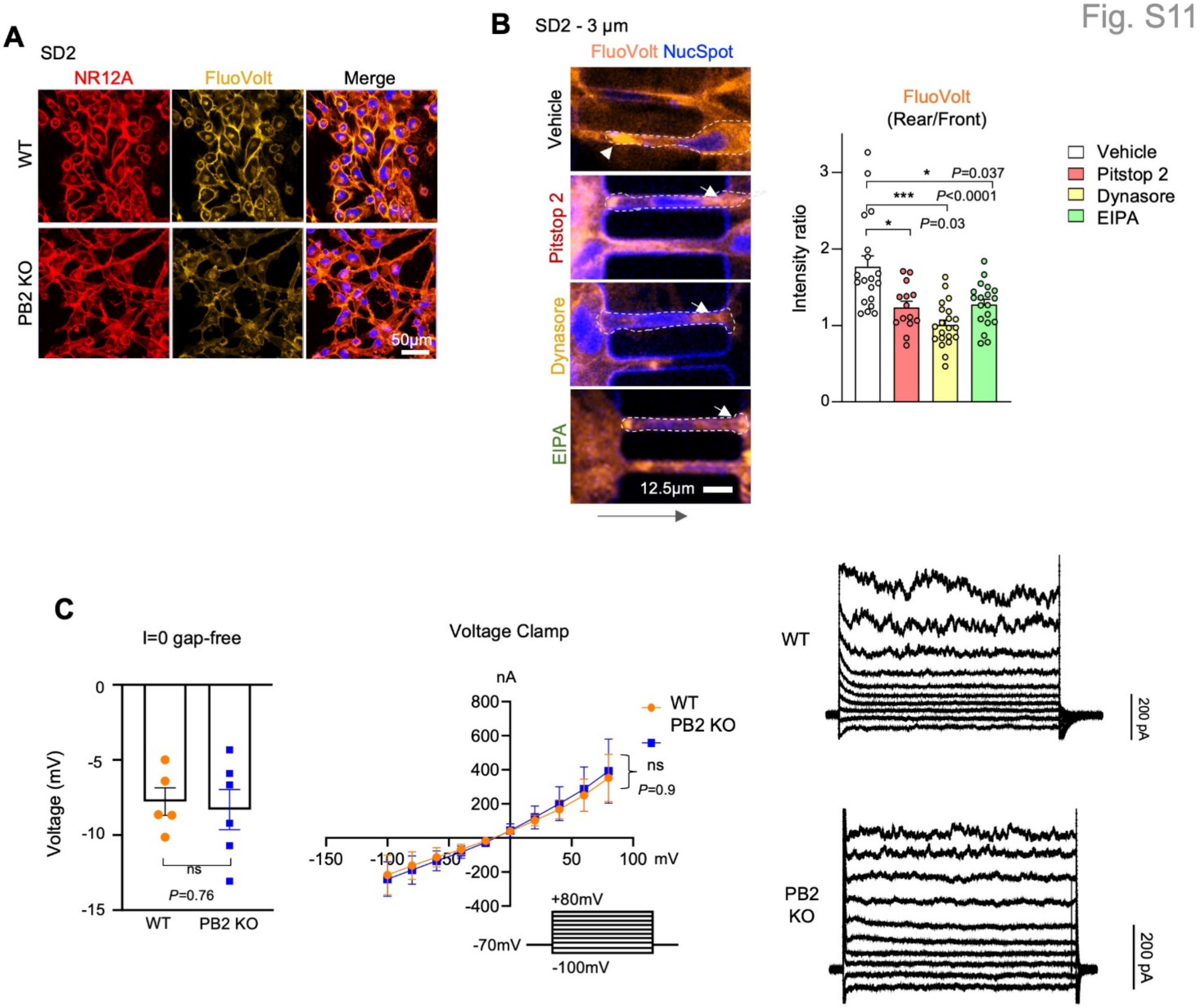


**Figure S11. Plexin-B2 affects the membrane electric field, but not transmembrane potential, related to Figure 6.**

(A) Live-cell imaging shows comparable Nile red (NR12) fluorescence intensity (solvatochromic plasma membrane-targeting dye), but reduced FluoVolt fluorescence intensity in PB2 KO compared to WT GSCs.

(B) Still images from videography show higher FluoVolt fluorescence signals at the rear zone (arrowhead) of WT GSCs traversing micro tunnel, a pattern disrupted by endocytosis inhibitors leading to FluoVolt fluorescence signals also appearing at cell front (arrows). Dashed lines delineate cell boundary. Long arrows denote migration direction. Right, bar graphs show the ratio of FluoVolt intensity at rear vs. front during confined migration. n=14-21 cells for each condition. Kruskal–Wallis test followed by Dunn's multiple comparisons test. Data represent mean ± SEM.

(C) Whole-cell patch-clamp recordings of SD2 GSCs. Left, currents were clamped at 0 A and bar graphs show comparable transmembrane potentials of WT (n = 5) and PB2 KO (n = 6) cells. Two-sided unpaired t-test. Middle, I/V curves reveal similar membrane conductance of WT (n=4) and PB2 KO (n=5) cells. The voltage step is shown below. Right, representative whole cell current traces from WT and PB2 KO cells. Cells were clamped in 20 mV steps relative to holding potential (- 70 mV), ranging from −100 to +80 mV. Two-way ANOVA followed by Bonferroni's post hoc test.

Fig. S12

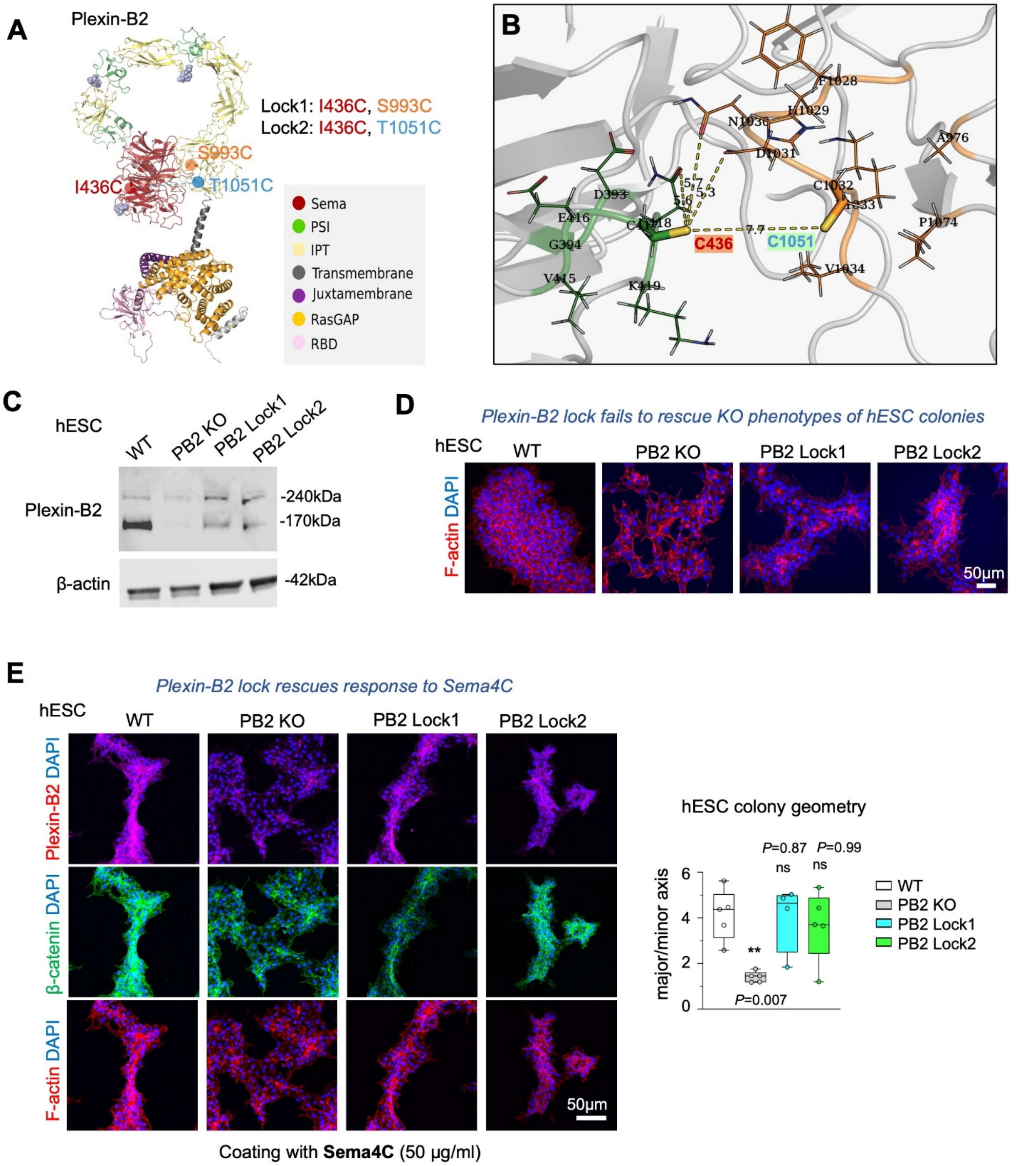


**Figure S12. Locked ring mutations of Plexin-B2 affect cytoskeleton organization, but not responsiveness to semaphorin, related to Figure 7.**

(A) 3D structural modeling of human Plexin B2, with domains coded by colors. Predicted glycosylations of residues N108, N509 and N714 are depicted as light blue spheres. Image visualization made with Pymol (version 2.3). The locations of two pairs of mutations: Lock1 (I436C and S993C) and Lock2 (I436C and T1051C) are predicted to form intramolecular disulfide bridges that lock the ring structure.

(B) Predicted intramolecular disulfide bond interaction of human Plexin B2 with mutations I436C and T1051C (Lock2). The predicted intramolecular disulfide bridges between C436 with C1051 locks the ring structure (note that the amino acid positions in the figure is shifted, as the signal peptide is excluded). Dashed yellow lines represent the distance between atoms, given in angstroms. Image visualization made with Pymol (version 2.3).

(C) Western blot shows absence of mature Plexin-B2 (170 kDa) in PB2 KO hESCs (H9 cell line) compared to wild-type cells, and re-expression of PB2 lock mutants. β-actin serves as a loading control.

(D) IF images of hESC colonies stained for F-actin (Alexa 568 phalloidin) and DAPI show differences in cell organization, colony geometry, and actin network between WT and PB2 KO, which were not rescued by PB2 Lock1 and Lock2 mutants.

(E) IF images of hESC colonies in culture dishes coated with Sema4C-Fc (50 µg/ml) mixed with Matrigel and stained for Plexin-B2, β-catenin, and F-actin. DAPI for nuclear staining. Sema4C affected the colony geometry of WT but not PB2 KO cells, and the insensitive to Sema4C was rescued by PB2 Lock1 and Lock2. n=5 colonies per group. One-way ANOVA followed by Dunnett's multiple comparisons to compare treatments within the groups.

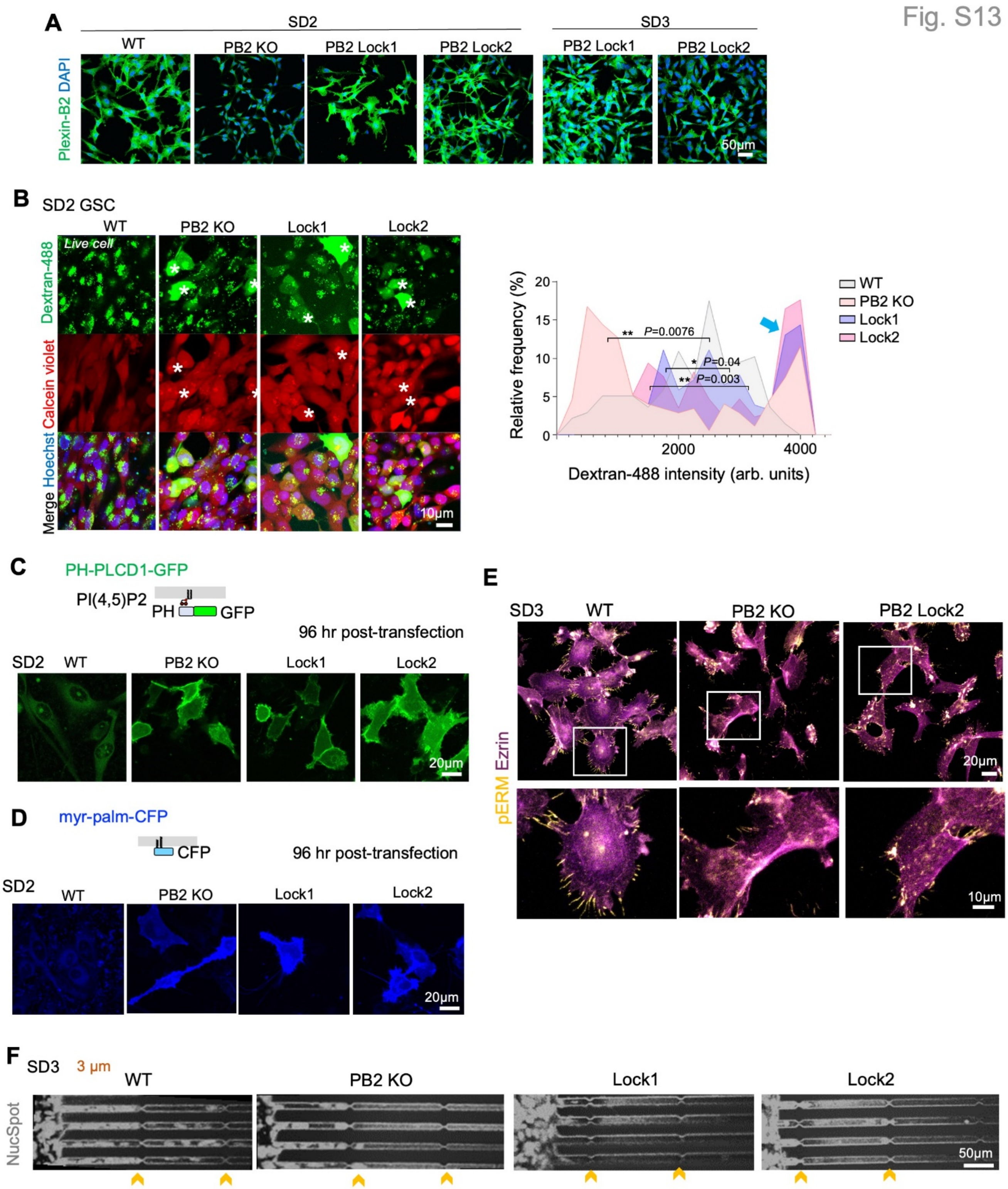


**Figure S13. Plexin-B2 extracellular locked ring mutation in glioma stem cells, related to Figure 7.**

(A) IF images show absence of Plexin-B2 in PB2 KO GSCs compared to WT SD2 and re-expression of PB2 Lock1 and Lock2 mutants in both SD2 and SD3. DAPI for nuclear staining. Also see WB shown in Fig. S5B.

(B) Live-cell confocal fluorescence images show diffuse cytoplastic dextran-Alexa488 (asterisks) in PB2 KO cells and both lock mutants. Calcein violet-AM dye was used to indicate live cells, Hoechst for cell nucleus. Histograms show bimodal distribution of dextran-Alexa 488 fluorescent intensities for PB2 KO and both lock mutants (blue arrow), reflective of reduced endocytosis and higher cytoplasmic dextran permeability. n=137 cells for WT, n=173 for PB2 KO, n=170 for Lock1, and n=153 for Lock2. Kruskal–Wallis test followed by Dunn's multiple comparisons test.

(C) Top, diagram of the PIP2 probe PH(PLCD1)-GFP. Bottom, live-cell imaging at 96 hr post transfection shows internalization of the PIP2 probe in WT GSCs in contrast to the membrane retention in PB2 KO and Lock mutants.

(D) Top, schematic of myr-palm-CFP attached to inner membrane leaflet. Bottom, live cell imaging at 96 hr after transfection shows membrane retention of the probe in PB2 KO and of both lock mutants, but not WT GSCs.

(E) Confocal IF images of different GSCs stained for pERM (phospho-Ezrin/Radixin/Moesin) and total Ezrin. Enlarged images of boxed areas are shown below. Note high pERM signals at tip of processes of WT cells compared to PB2 KO cells, and failure of rescue by PB2 Lock2 mutant form.

(F) Still images at the end of 17 hr videography show compromised confined migration of PB2 KO and both lock mutants through microchannels with 3 μm constrictions as compared to WT. NucSpot for live cell nuclei staining.

**Supplemental video legends**

**Movie S1. Live-cell imaging of GSCs migration through 3 and 8 µm constrictions (related to Figure 1).**

SD2 wild-type (WT) glioma stem cells (GSCs) migrating through microchannels containing 3 or 8 µm constrictions, imaged every 5 min over approximately 21 hr. NucSpot for nucleus staining. Note that in contrast to 8 µm constrictions, GSCs passing through the 3 µm constrictions displayed higher speed and higher momentum moving forward.

**Movie S2. Live-cell imaging of SD2 and SD3 glioma stem cells migrating through 3 µm constrictions (related to Figure 1).**

SD2 and SD3 wild-type (WT) GSCs migrating through microchannels with 3 µm constrictions, imaged every 5 min over approximately 16 hr. NucSpot for nucleus staining. Note that SD3 traversed the 3 µm constrictions at a higher velocity than SD2.

**Movie S3. Live-cell imaging of human neuroprogenitor cells and SD3 glioma stem cells migrating through 3 µm constrictions (related to Figure 1).**

Human neuroprogenitor cells (hNPCs) derived from human pluripotent stem cells and SD3 wild-type GSCs migrating through microchannels with 3 µm constrictions, imaged every 5 min over approximately 16 hr. NucSpot for nucleus staining. Note that hNPCs displayed lower migratory capacity to negotiate 3 µm microchannels than GSCs.

**Movie S4. Live-cell imaging of GSCs treated with endocytosis inhibitors (related to Figure 1).**

SD2 wild-type (WT) GSCs migrating through microchannels with 3 µm constrictions, treated with endocytosis inhibitors (PitStop2, Dynasore, or EIPA) or vehicle and imaged every 5 min over approximately 17 hr. NucSpot for nucleus staining.

**Movie S5. Plexin-B2 deficient GSCs show long tethers during migration in 2D cell culture (related to Figure 2).**

The videos show SD2 GSCs with CRISPR/Cas9-mediated *PLXNB2* knockout (PB2 KO) migrating in laminin coated 2D cell culture. Solvatochromic plasma membrane dye Nile red NR12A (red) and NucSpot (blue) were used for staining cells, and imaged every 5 min over 13 hrs. No obvious overall motility impairment was observed, but PB2 KO cells exhibited fluidic cell membranes that often extended into long tethers, in contrast to wild-type (WT) GSCs.

**Movie S6. Live-cell imaging of SD2 WT and Plexin-B2 KO GSCs migrating through 3 µm constrictions (related to Figure 4).**

The videos show SD2 wild-type (WT) and CRISPR/Cas9-mediated *PLXNB2* knockout (PB2 KO) GSCs migrating through microchannels with 3 µm constrictions, imaged every 5 min over approximately 14 hr. SPY555 for F-actin (red), MemGlow (green) for cell membranes, and NucSpot (blue) for nuclei.

**Movie S7. Live-cell imaging of SD3 WT and Plexin-B2 KO GSCs migrating through 3 µm constrictions (related to Figure 4).**

The videos show SD3 wild-type (WT) and CRISPR/Cas9-mediated *PLXNB2* knockout (PB2 KO) GSCs migrating through microchannels with 3 µm constrictions, imaged every 5 min over approximately 15 hr. SPY555 for F-actin (red), MemGlow (green) for cell membranes, and NucSpot (blue) for nuclei.

**Movie S8. Plexin-B2 deficient GSCs show long tethers during migration through microchannels (related to Figure 4).**

The videos show SD2 wild-type (WT) and CRISPR/Cas9-mediated *PLXNB2* knockout (PB2 KO) GSCs migrating through microchannels with 8 µm constrictions, imaged every 5 min over approximately 14 hr. SPY555 for F-actin (red) and NucSpot (blue) for nuclei. Note the long tethers containing F-actin between PB2 KO GSCs attempting to dissociate from neighbor cells.

**Movie S9. Migration through long constricted tunnels requires actin polarization and endocytosis (related to Figure 4).**

The videos show SD3 wild-type (WT) GSCs migrating through long micro tunnels of 3 µm width and displaying actin assembly at rear and endocytosis at the front, both reduced in PB2 KO cells that were stalled and unable to squeeze through long micro tunnels. SPY555 (red) for F-actin, MemGlow (green) for cell membranes, and NucSpot (blue) for nuclei. Cells were imaged every 5 min over approximately 15 hr.

**Movie S10. Molecular dynamics simulation of cortical contractility and membrane tension determining the rate of endocytosis in confined migration (related to Figure 5).**

The videos show increasing confinement of two cells, modeled with a coarse-grained bead-spring model. The left video represents membrane-cytoskeleton energy setting of $U_3$=15, mimicking high membrane-cytoskeleton energy, and the right represents setting of $U_3$=0.03, mimicking low membrane-cytoskeleton energy. When the confinement reaches 0.6 of cell diameter, for $U_3$=0.03, the nucleus is loose within the cytoplasm and there is no formation of involution. In contrast, for $U_3$=15, the involution is generated by the high force in the heads of the actin filaments during the constriction. The simulations were performed using membrane elastic constant $k_{00}$=500. The beads represent actin filaments (white), actin filament head (yellow), nuclear envelope (green), cell membrane (red), and constrictions are represented by two opposite beads (light pink).

**Movie S11. Molecular dynamics simulation of cortical contractility and membrane tension determining cell movement rate under confinement (related to Figure 5).**

The videos show invasiveness of three cells under confinement. The beads represent actin filaments (red), actin filament head (black), nuclear envelope (green), cell membrane (yellow), and constrictions are represented by two opposite lines. When the confinement reaches 0.6 of the cell diameters, the intermediate values of membrane-cytoskeleton energy and cell membrane energy ($U_3$=6, $\kappa_{00}$=100) promoted sustained unidirectional confined migration. For low values ($U_3$=0.03, $\kappa_{00}$=10) the cell moved for a short period in one direction and suddenly changed directionality. For high values ($U_3$=30, $\kappa_{00}$=5000), the cell displayed a hypercontractile state with reduced displacement.

**Movie S12. Migration through constricting tunnels involves high FluoVolt signals at rear zone of the GSCs (related to Figure 6).**

The videos show SD2 wild-type (WT) GSCs migrating through micro tunnels of 3 μm width, displaying high FluoVolt signals at rear end (i.e., less negative charge), in contrast to PB2 KO cells that showed reduced FluoVolt signals and were stalled and unable to squeeze through tunnels. FluoVolt (orange) is a voltage-sensitive fluorescent membrane dye and NucSpot (blue) was used to stain nuclei. Cells were imaged every 5 min over approximately 17 hr.

**Movie S13. Migration through constricting tunnels involves calcium signaling in GSC (related to Figure 6).**

The videos show SD2 wild-type (WT) GSCs migrating through chambers and narrow tunnels of 3 μm width, with or without treatment with the calcium chelator BAPTA-AM. BAPTA treated GSCs showed reduced FluoVolt signals and were stalled in chambers and unable to squeeze through tunnels. FluoVolt (orange) is a voltage-sensitive fluorescent membrane dye and NucSpot (blue) was used to stain nuclei. Cells were imaged every 5 min over approximately 17 hr.
